# Genetic Insights of Image-Based Traits: Analysis Pipeline for AI-based Phenotyping, Combined-GWAS, and Federated Learning with Application to the Human Face

**DOI:** 10.1101/2025.08.03.668352

**Authors:** Xianjing Liu, Ziyi Xiong, Fan Liu, Tamar Nijsten, Eppo B. Wolvius, Manfred Kayser, Gennady V. Roshchupkin

**Author notes:** These authors contributed equally to this work (Xianjing Liu, Ziyi Xiong). These authors contributed equally to this work (Manfred Kayser, Gennady V. Roshchupkin). **Corresponding author:** Gennady Roshchupkin. **IRB approval status:** The Rotterdam Study has been approved by the Erasmus MC Medical Ethical Committee (MEC-02-1015), and by the Dutch Ministry of Health, Welfare and Sport (Population Screening Act, reference 3295110-1021635-PG). The Generation R Study was approved by the Erasmus MC Medical Ethics Committee (MEC 198.782/2001/31).

## Abstract

Biological images capture rich morphological information and understanding their genetic basis is crucial for elucidating underlying molecular mechanisms, relevant in biomedical, evolutionary, and forensic research and applications. However, progress in understanding the genetic basis of image-based complex traits is hindered by key limitations in the current methodology, regarding both, the degree of which the large image complexity is captured with the phenotyping methods and how the analysis methods deal with the underlying large genetic complexity. Here, we present a robust, scalable, privacy-preserving computer pipeline for analysing image-based complex traits and unveiling their genetic basis by integrating (i) AI-based phenotyping for large-scale extraction of image-based traits across cohorts in a federated learning framework, without sharing individual images; (ii) Combined-GWAS (C-GWAS) for identifying genetic variants underlying the numerous AI-derived image-based traits; and (iii) Explainable AI (XAI)-based image visualization of the identified genetic association effects. In its first application to the example of the human face, we analysed 3D facial images and genomic data from two European cohorts (N=7,309), extracted 200 AI-derived facial traits, identified 43 significantly face associated genetic loci, including 12 novel ones, and replicated 70% of them in an independent European dataset (N=8,246). XAI-based visualization of the identified genetic effects shows the involvement of many of these loci in different parts of the face. Our study provides a privacy-aware and extensible pipeline for investigating the genetic basis of image-based complex traits implemented in a computationally efficient python package available at GitLab and its first application yielded new insights into the genetic architecture of facial shape variation.

## Introduction

Biomedical imaging—ranging from 3D scans of full bodies, certain body parts or particular organs, to histopathology slides—captures complex structural and textural patterns of the human body. Understand the genetic basis of image-based complex traits is relevant in many areas of life sciences including medicine. Yet, defining phenotypes that capture the full spectrum of a biological image is challenging with existing image analysis methodology. Several strategies have been proposed to tackle the challenges when it comes to analysing image-based complex traits and unveiling their genetic basis, such as utilizing using each pixel or voxel within an image e.g. brain magnetic resonance imaging (MRI)^11^ or prior-defined measurements e.g. as landmarks from facial images^9^ or the region-of-interest volume from brain MRI scans^10^ and using them as phenotypes for genome-wide association studies (GWASs) to unveil their genetic basis. Although these strategies yield commendable outcomes when applied to large sample size^9,10,11,12^, the real potential lies in seeking solutions that uncover more comprehensive, data-driven image-based phenotypes that move beyond a priori definitions, and better reflect the underlying biological variation.

Recent breakthroughs in artificial intelligence (AI) models have demonstrated promising data-driven capabilities for analysing images by reducing high-dimensional images into low-dimensional representations, i.e., AI-derived traits, while preserving all essential biological information the image contains. This approach has proven fruitful in various GWAS contexts, including unsupervised autoencoder-derived traits from 3D brain MRI^14^, 3D cardiac mesh images^15^, and 3D facial mesh images^80^, as well as traits learned via self-supervised contrastive learning^16^ and transfer learning^75^ from 2D retinal fundus images.

However, until now, all GWASs of AI-derived traits have applied the classical singled trait approach—testing each AIdderived trait separately and in isolation from another of the same image. Yet, these image-based AI-derived traits often are inherently correlated with each other, as result of underlying biological such as genetic correlations. Classical single-trait GWAS has limited ability to detect genetic loci jointly influencing these genetically correlated traits with sufficient statistical power. To effectively combine single-trait GWASs of large numbers of correlated (or uncorrelated) traits, such as those derived from bioimages, some of us recently introduced a combined genome-wide association study approach named Combined-GWAS (C-GWAS)^6^. The largely increased power of C-GWAS relative to meta-analysis of single-traits GWASs has been demonstrated in both, simulated data as well as real data applications using landmark-based phenotypes of the human ear^76^ and the human face^6^. Yet, C-GWAS has not been applied to image-based AI-derived traits.

A challenge in AI-based phenotyping of bioimages is the standardization of AI-derived traits across cohorts when they belong to different institutions. Genetic studies of complex traits caused by many genes with small effect sizes require very large sample sizes, often necessitating the combination of data from different cohorts of multiple institutions. For traditional traits, GWAS summary statistics derived from standardized measurements can be shared between institutions without exchanging individual image data. However, AI-derived traits are inherently shaped by cohort-specific training data distributions. Consequently, AI models trained separately in different cohorts may produce AI-derived traits that are not directly comparable between the cohorts, providing challenges for successful GWAS meta-analysis. Sharing bioimages of different cohorts between different institution, as needed to train a unified model, typically is constrained by restrictive privacy regulations. Federated learning addresses this challenge by enabling multiple sites to collaboratively train a single AI model^56^, resulting in aligned AI-derived traits across cohorts—without sharing individual image data.

Here, we present a robust, scalable, privacy-preserving analysis pipeline for analysing image-based complex traits and exploring their genetic basis, which integrates (i) Federated AI-based phenotyping of images for large-scale extraction of image-based traits across cohorts without sharing individual images; (ii) Combined-GWAS (C-GWAS) for identifying genetic variants underlying the numerous AI-derived image-based traits; and (iii) Explainable AI (XAI)-based visualization of the identified genetic association effects on the image. To empirically exemplify its practical suitability, we applied our combined image and genetic analysis pipeline to the human face, by using 3D digital facial images and genome-wide data of thousands of Europeans from several cohorts.

## Results

We developed an integrated computer pipeline for analysing biological images and obtaining genetic insights into image-based complex traits, which combines i) data-driven AI-based phenotyping to cover image complexity via a large number of AI-derived traits, implemented within a federated learning framework without sharing individual images across cohorts, ii) combined genome-wide association studies (C-GWAS) from a large number of image-based AI-derived traits, iii) Explainable AI (XAI)-based visualization of the identified genetic association effects on the full image. We applied this framework to the prominent example of the human face (Figure 1).

**Figure 1:**
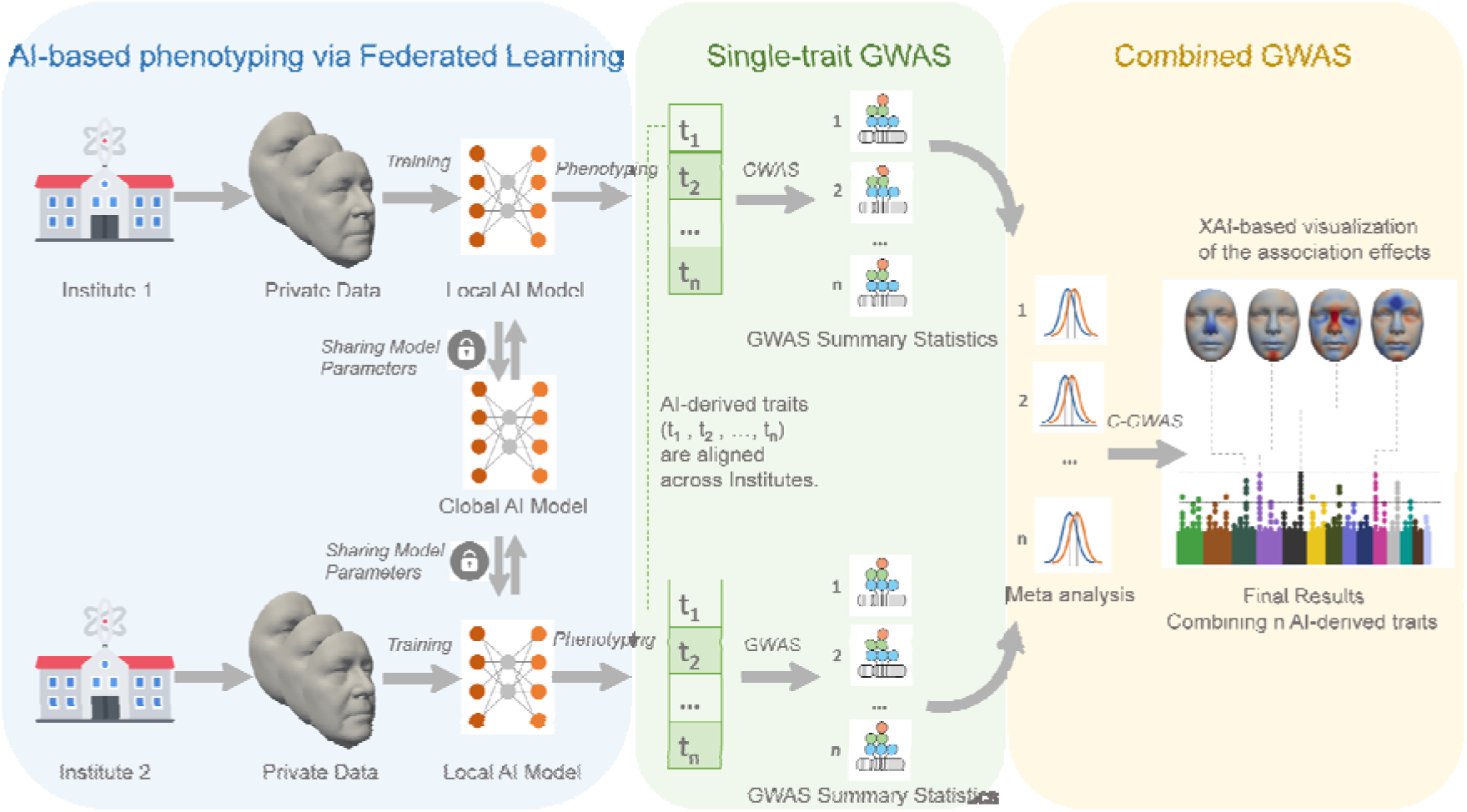
Computer pipeline to analyse complex biological images and obtain genetic insights into image-based complex traits using the human face as an example. The analysis pipeline consists of three blocks depicted with different underlaying colour shades: 1) AI models were trained via federated learning techniques without inter-institutional sharing individual-level image data and extraction of numerous AI-derived traits from the full image (light blue); 2) single-trait GWAS were performed for each AI-derived trait within each cohort separately (light green); 3) meta-analysis of all single trait GWAS summary statistic data from all AI-derived traits and all cohorts and C-GWAS for combing all AI-derived traits from all cohorts with explainable AI (XAI)-based visualization of the revealed genetic association effects on the full image (light yellow).

### Comparison between federating learning and centralized training strategies

AI-based phenotyping requires model-based learning. In traditional CT, images need to be shared between study sites for merging them into one single training session. In practise, due to privacy regulations, inter-institutional image sharing often is not possible, especially for privacy-sensitive images such as facial images. Applying FL to image-based analysis avoids the need of inter-institutional individual image sharing. Figure 2 illustrates the conceptual difference between the FL and CT training strategies. To evaluate whether FL performs comparably to CT, we compared the image reconstruction performance of our AI-based facial phenotyping model trained using FL and CT (Table 1).

**Figure 2:**
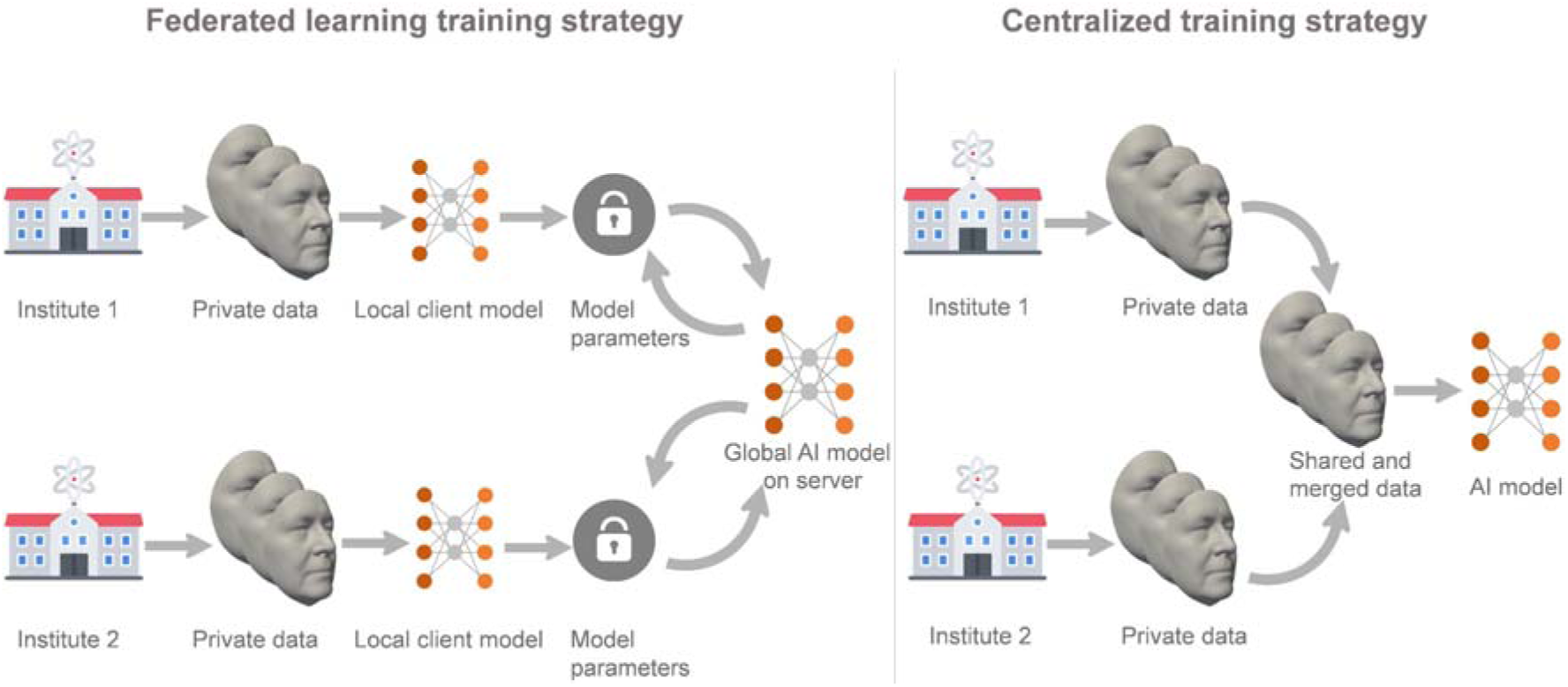
Overview of the conceptual difference between federated learning (FL) training strategy and centralized training (CT) strategy for bioimages using the example of the human face.

**Table 1:** Mean reconstruction error of FL and CT strategies.

|  | CT (mm) |  | FL (mm) |  |
| --- | --- | --- | --- | --- |
|  | Train (80%) | Test (20%) | Train (80%) | Test (20%) |
| **RS** (N=3,995) | 0.229 ± 0.031 | 0.232 ± 0.032 | 0.232 ± 0.031 | 0.235 ± 0.033 |
| **GenR** (N=3,314) | 0.223 ± 0.017 | 0.227 ± 0.019 | 0.226 ± 0.018 | 0.229 ± 0.020 |
| **RS + GenR** (N=7,309) | 0.226 ± 0.026 | 0.230 ± 0.027 | 0.229 ± 0.026 | 0.233 ± 0.028 |

CT: model was trained by a centralized dataset with merged images from both cohorts RS and GenR, FL: model was trained via FL settings without creating a centralized dataset across both cohorts.

We found that our proposed FL training strategy (combined cohort mean test error 0.230 ± 0.027 mm) demonstrated comparable performance without individual image sharing relative to the traditional CT strategy (combined cohort mean test error 0.233 ± 0.028 mm) that requires individual image sharing (Table 1). Across both training strategies, reconstruction errors on the training and test sets were highly consistent, indicating stable generalization performance without evidence of substantial overfitting. Figure S1 shows details about the reconstruction error visualized as facial heatmaps for an individual from the elderly cohort (RS) and an individual from the children cohort (GenR) as representative examples. These examples illustrate that the overall facial morphology is well preserved across cohorts with distinct age distributions. Our results indicate that FL is an effective strategy for image-based AI model training, at least when it comes to the face example used here, thereby bypassing the need of sharing individual image data between different institutions. Notably, larger reconstruction errors were found in sharp facial areas such as eyes and nasolabial folds. Correspondingly, slightly higher reconstruction errors were found in the RS cohort compared with GenR.

In addition, Figure S9 provides qualitative visual evidence that the AI-derived traits learned by the FL model are well aligned between the two cohorts. Using facial heatmap visualizations, corresponding traits show consistent facial patterns between RS and GenR. This further supports that FL enables consistent cross-cohort phenotyping without sharing individual-level image data.

### C-GWAS and replication outcomes

After performing single trait GWASs on all 200 AI-derived facial traits in both European cohorts separately, and meta-analysis of both cohorts (N=7,309), we carried out C-GWAS to search for genetic associations in both cohorts in a combined way. To visualize the phenotypic effects of the obtained genetic associations, we developed an explainable AI (XAI)-based approach that maps C-GWAS-derived SNP effects back onto interpretable facial morphology patterns (Figure 3). Implementation details of this approach are described in the Methods section.

**Figure 3:**
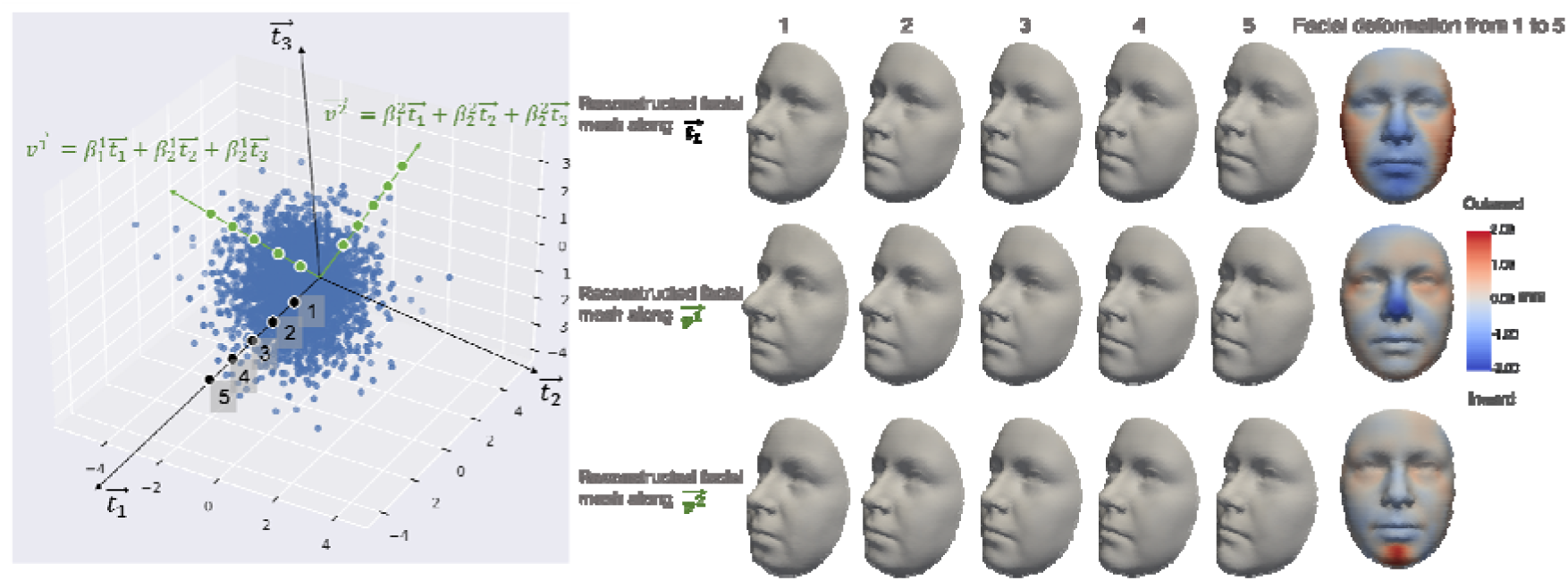
Explainable AI (XAI)-based facial visualisation of genetic association effects. Facial images were reconstructed by sampling along vector directions in the AI-derived trait space (n = 3 dimensions for demonstration). Axis,, …, are basis vectors representing n AI-derived facial traits; β^i^_1_, β^i^_2,_ …, β^i^_n_ are the effect size of SNP^i^ for the n traits obtained from single-trait GWAS. Vectors and represent the composite genetic effect directions for SNP^1^ and SNP^2^ derived from the C-GWAS results. When sampling along a vector, a series of facial images were reconstructed from the decoder, and outward changes in the reconstructed faces (with respect to the geometric center of the head) were shown as red regions in the heatmap, while blue regions indicate inward changes. Each blue dot in the diagram is a latent data point representing the low-dimensional representation of an input facial image in the trait space.

Based on C-GWAS in both European cohorts combined, we identified 43 genetic loci demonstrating SNPs with genome-wide statistically significant facial associations (Figure 4, Figure S6, Table S1 for more details). Via AI-derived facial heatmaps, we demonstrate the phenotypic effects of the obtained significant associations (Figure 4). An inspection of these 43 loci in all available resources, i.e., a comprehensive review of the previous human face genetic literature (Table S2) and a recently published large-scale face C-GWAS in Europeans^49^, revealed that 12 (28%) of these loci were not previously reported as face loci, marking them as novel genetic face loci we identified in this study. Regional association plots for these 12 novel loci are provided in Figure S10.

**Figure 4:**
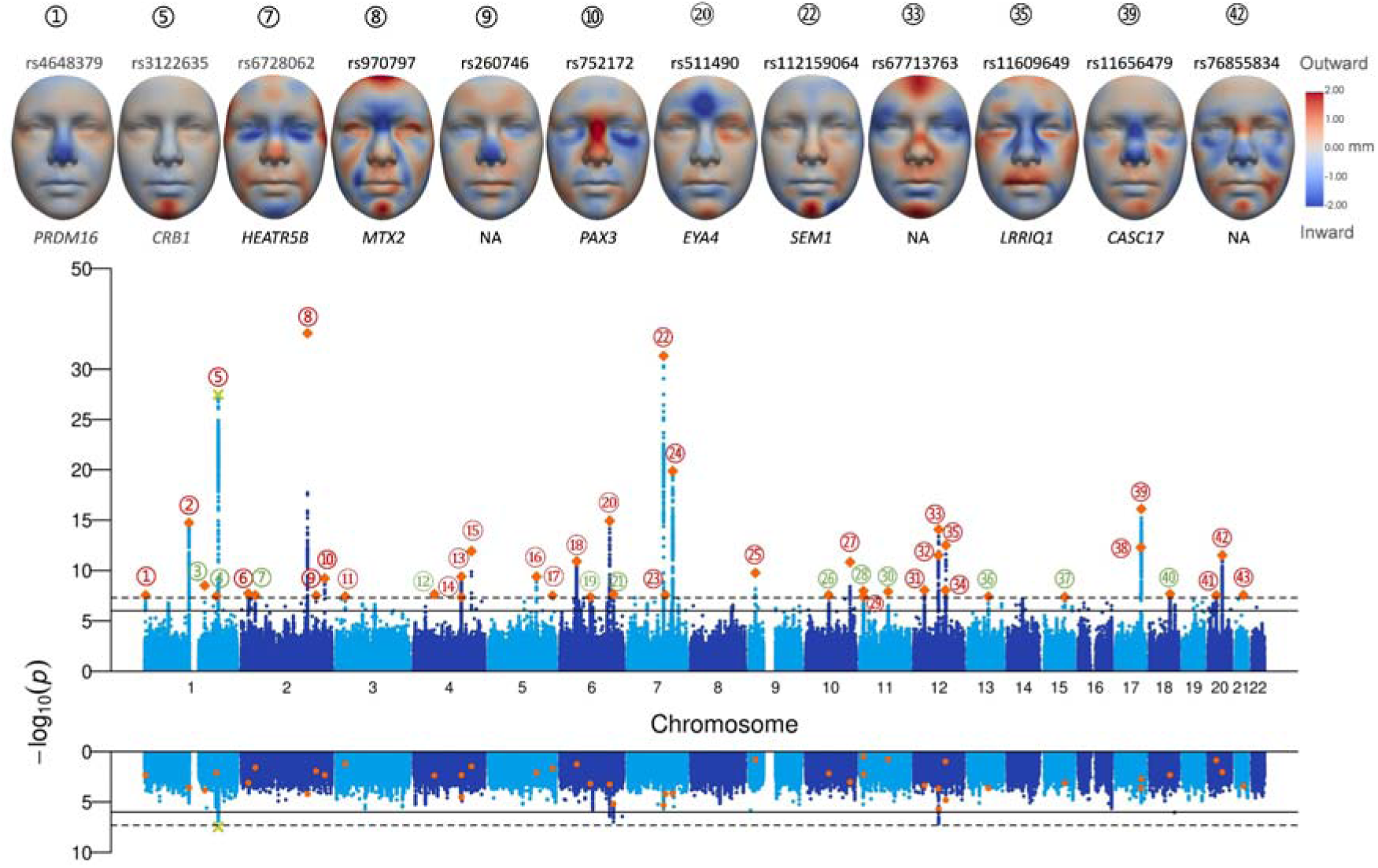
Manhattan plot of C-GWAS outcomes (upper part) from 200 AI-derived facial traits in two European cohorts combined (N=7,309) and AI-based facial visualization of the association effects of the C-GWAS-derived top SNPs. Green numbers with circles denote novel face loci, while red numbers denote previously known face loci that were confirmed here. Lower part shows results from single-trait GWAS meta-analysis of both cohorts; green-crosses denote SNPs that were significant in both, single-trait GWAS and C-GWAS; red-dots denote SNPs that were only significant in C-GWAS. Red color in the heatmap means outward facial effects w.r.t the geometric center of the head, while blue means inward facial effects.

The AI-derived facial heatmap visualization of the genetic association effects of the 12 novel loci demonstrated that they are all linked with several facial traits involving multiple facial regions (Figure 5). This finding underlines the ability of C-GWAS in identifying genetic loci with pleotropic effects.

**Figure 5:**
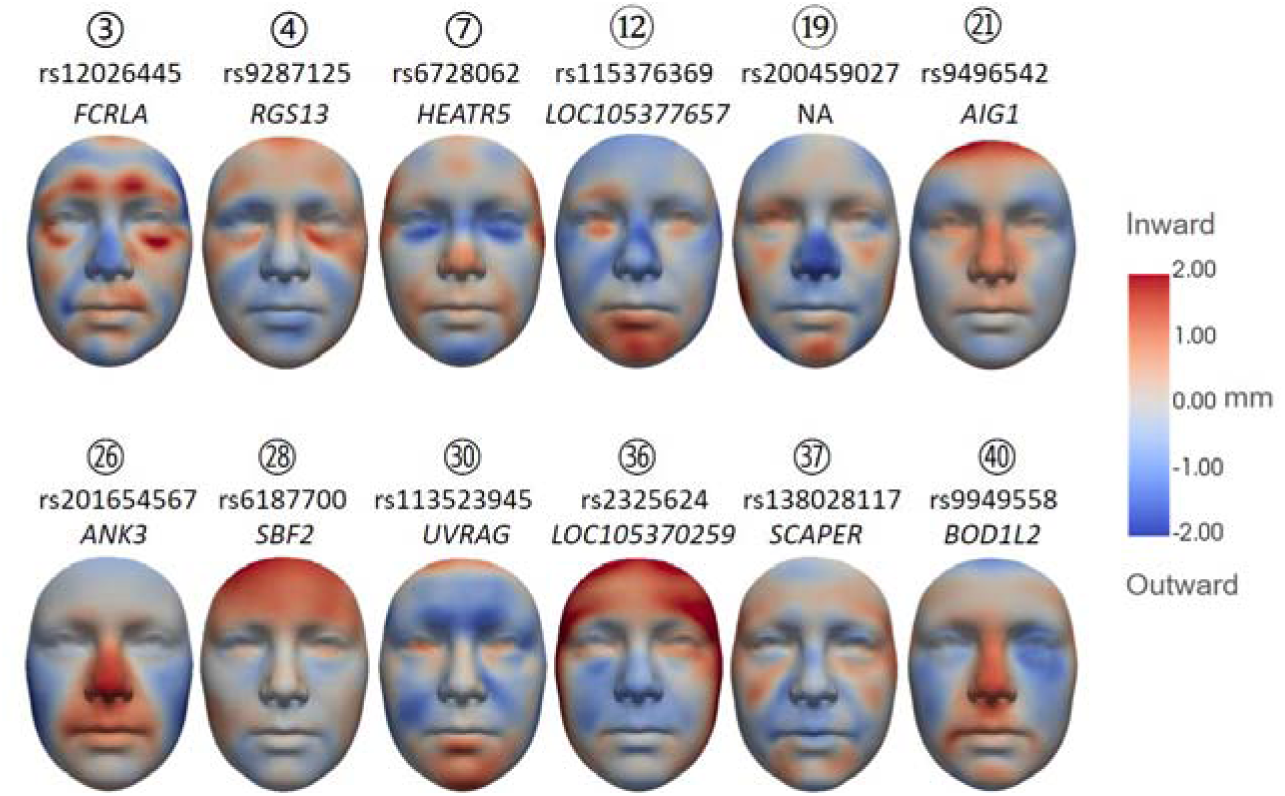
AI-derived facial heatmaps visualizing the association effects of the 12 novel genetic face loci highlighted by the C-GWAS of both European cohorts combined. Red color means outward effects w.r.t the geometric center of the head, while blue means inward effects.

According to annotation information of FUMA^77^, the 12 novel loci are additionally linked to other traits, such as QRS duration (⍰)^54^, brain morphology (⍰ □)^57^, diabetic maculopathy (⍰)^58^, blood cells (⍰ □)^59^, blood pressure measurement (□ □)^60,61^, total cholesterol measurement(□)^62^, and Alzheimer disease (□)^63^.

In the replication analysis using independent European data (N=8,246)^5^, 30 (70%) of the 43 loci, including two novel loci, rs9496542 (*AIG1/LOC124901231*) and rs113523945 (*UVRAG*), demonstrated statistically significant associations (i.e., p < 0.05), providing confirmatory evidence (Table S1). Figure S3 shows the comparison of facial heatmaps between our findings from the discovery analysis and the replication analysis, providing further evidence that the identified loci exhibit consistent phenotypic patterns in both analyses.

### Comparison of C-GWAS outcomes between the two cohorts

Since both discovery cohorts include participants of largely different age ranges, we additionally performed C-GWAS of the 200 AI-derived traits separately for the children in GenR (N=3,314) and the elderly in RS (N=3,995) (Manhattan plot in Figure S2). With 21 identified genome-wide significant genetic loci in RS and 18 in GenR, we found about similar number of loci in the elderly cohort as in the children cohort; however, only four loci (*CRB1*; *MTX2; EYA4; SEM1*) overlapped between both cohorts.

We further focused on the changes of the statistical power before and after meta-analysis of both cohorts in C-GWAS. Among the 43 identified genetic loci (Table S1), 37 (86%) showed increased power after meta-analysis of both cohorts (Figure 6). For instance, four loci were genome-wide significant in both cohorts separately and reached stronger significance in the meta-analysis of both cohorts (Figure 6a) and four loci were genome-wide significant in one of the two cohorts and genome-wide significance in the meta-analysis of both cohorts (Figure 6b,c), and five loci were not genome-wide significant in either of the two cohorts, while they reached borderline or nominal significance in both cohorts, showed consistent patterns in facial heatmaps across both cohorts, and were genome-wide significant in the meta-analysis of both cohorts (Figure 6d). These findings highlight the benefits of increased sample size in enhancing statistical power through our proposed pipeline despite the different age ranges of the two cohorts.

**Figure 6:**
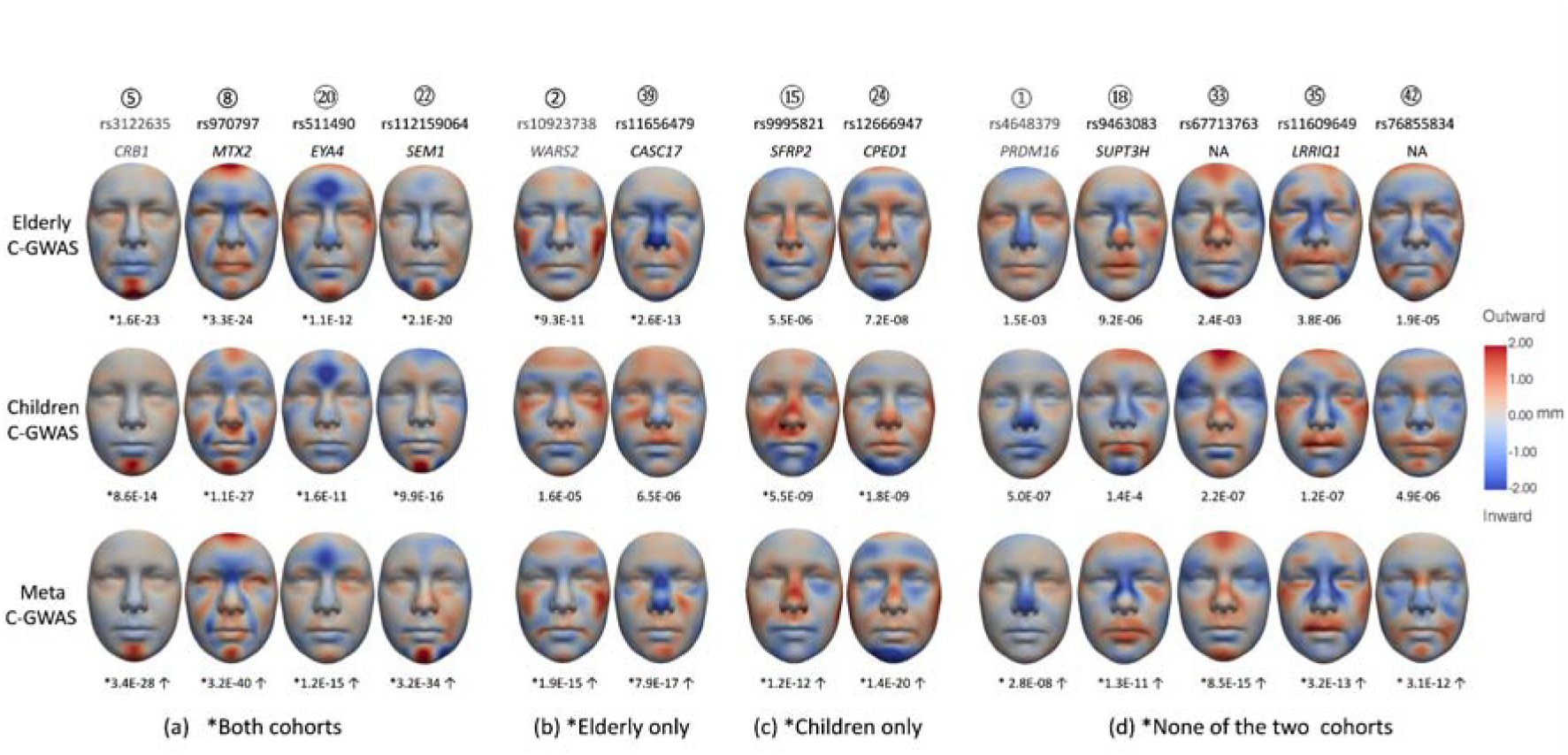
Impact of meta-analysis on loci detection power with C-GWAS. a) Loci genome-wide significant in the two separate C-GWASs in both cohorts; b) Loci genome-wide significant in the separate C-GWAS of the elderly RS cohort; c) Loci genome-wide significant in the separate C-GWAS of the children GenR cohort; d) Loci genome-wide significant in the meta-analysis C-GWAS of both cohorts but non-significant in the two separate C-GWASs of both cohorts. Red color in the heatmap means outward effects w.r.t the geometric center of the head, while blue means inward effects. Numbers with circles are index of the 43 identified genetic loci in Table S1.

However, we also found 5 loci that were genome-wide significant in one of the two cohorts, but not anymore after meta-analysis of both cohorts (Figure 7). The facial visualizations show that these 5 SNPs exhibit different patterns of genetic effects across the two cohorts, as maybe expected. Importantly, these differences do not reflect inconsistencies in AI-derived traits across cohorts, but rather cohort-specific genetic effects acting on a shared phenotypic space. These findings may reflect underlying cohort-specific effects, such as the large age range difference of the participants of both cohorts.

**Figure 7:**
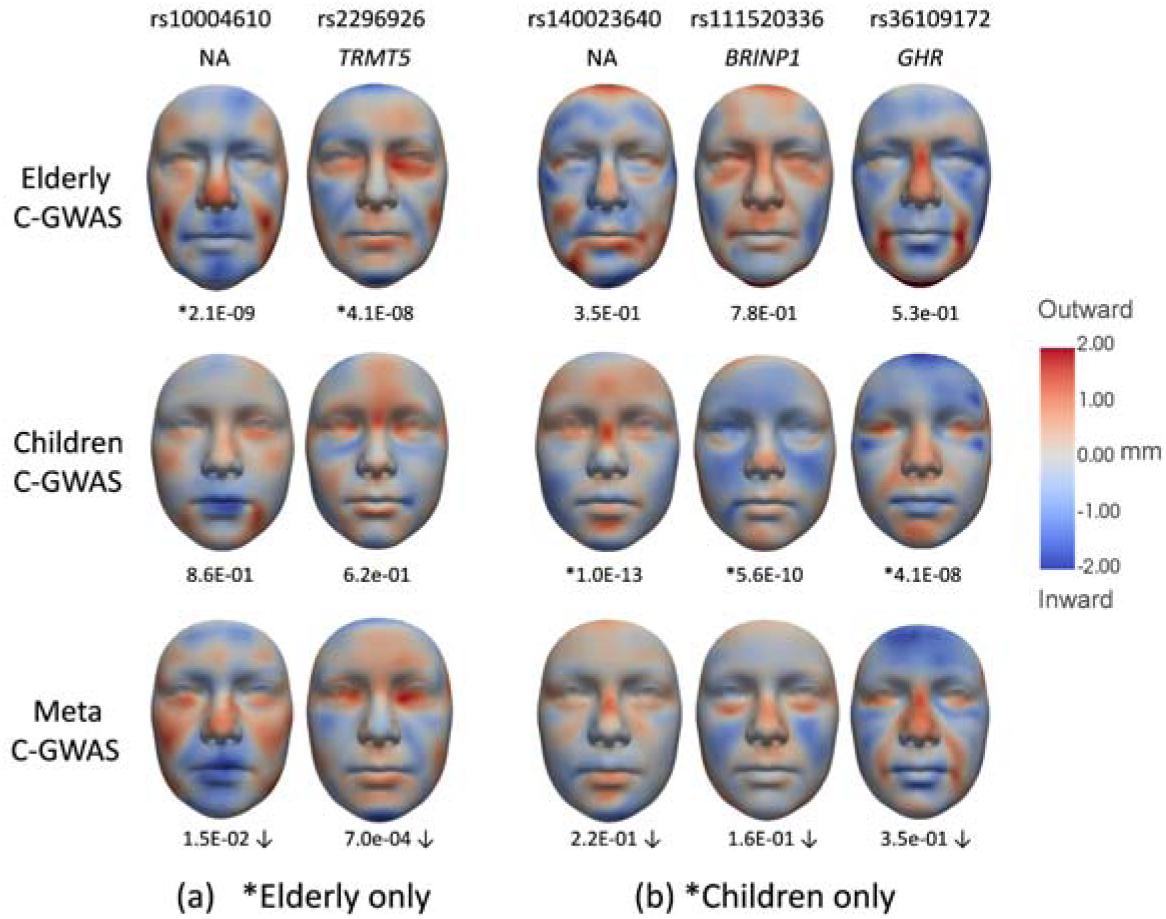
Genetic loci with genome-wide significance in separate C-GWASs in one of the two cohorts, but not in the C-GWAS of both cohorts combined. a) Genome-wide significant loci in the elderly RS cohort, but not in the children GenR cohort; b) Genome-wide significant loci in the children GenR cohort, but not in the elderly RS cohort. Red color in the heatmap means outward effects w.r.t the geometric center of the head, blue means inward effects.

### Heritability analysis outcome

LDSC-based heritability analysis on the single-trait GWAS summary statistics after meta-analysis of the two cohorts was performed and after false discovery rate (FDR) correction for multiple testing, revealed a total of 90 of the 200 (45%) AI-derived traits to be statistically significant with the h^2 range from 0.13 to 0.34 (Table S3). This finding is remarkable because only one genetic locus was identified with genome-wide significant association in the single-trait meta-analysis GWAS results (Figure 4). Next, we combined these FDR-significant traits and mapped them onto the face, revealing combined heritability effects ranging from 0.20 to 0.35. As evident from Figure 8, almost all facial regions are covered. Figures S4 and S5 show implementation details about merging multiple traits and mapping them back to the face.

**Figure 8:**
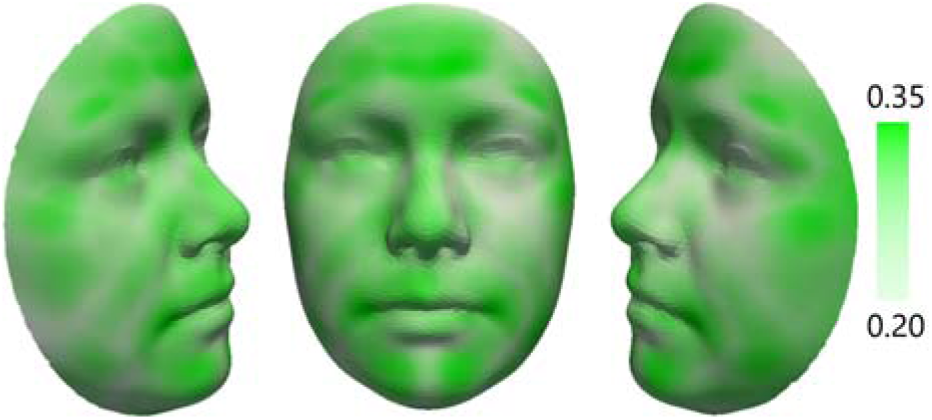
LDSC-based heritability of facial traits from meta-analysis of single-trait GWAS summary statistics based on the combination of 90 FDR-significant facial traits.

### Genetic correlation outcome

We first examined the phenotypic correlations between the 200 AI-derived facial traits (Figure S8). The distribution of pairwise correlations across the 200 traits followed an approximately Gaussian distribution with a mean of 0.000□±□ 0.084 (SD). Accordingly, about 95% of the correlations fell within 0.16 and 99.7% within 0.25, indicating overall low correlations among the AI-derived traits and suggesting that they capture largely distinct aspects of facial morphology.

A LDSC-based genetic correlation analysis between the 200 AI-derived facial traits based on single-trait GWAS summary statistics after meta-analysis of the two cohorts highlighted 21 (10.5%) of the 200 traits to be genetically correlated with at least one other trait based on FDR correction (α□ **=**□0.05) performed separately for each trait across its 199 pairwise tests. Among the significant pairs (Figure 9), absolute genetic correlations ranged from 0.56 to 0.92 (mean: 0.73), whereas the corresponding phenotypic correlations ranged from 0.05 to 0.67 (mean: 0.23). While correlations among AI-derived traits are expected given the absence of orthogonality constraints, the substantially higher genetic correlations compared with phenotypic correlations indicate shared genetic influences across multiple AI-derived traits, suggesting pleiotropic genetic effects.

**Figure 9:**
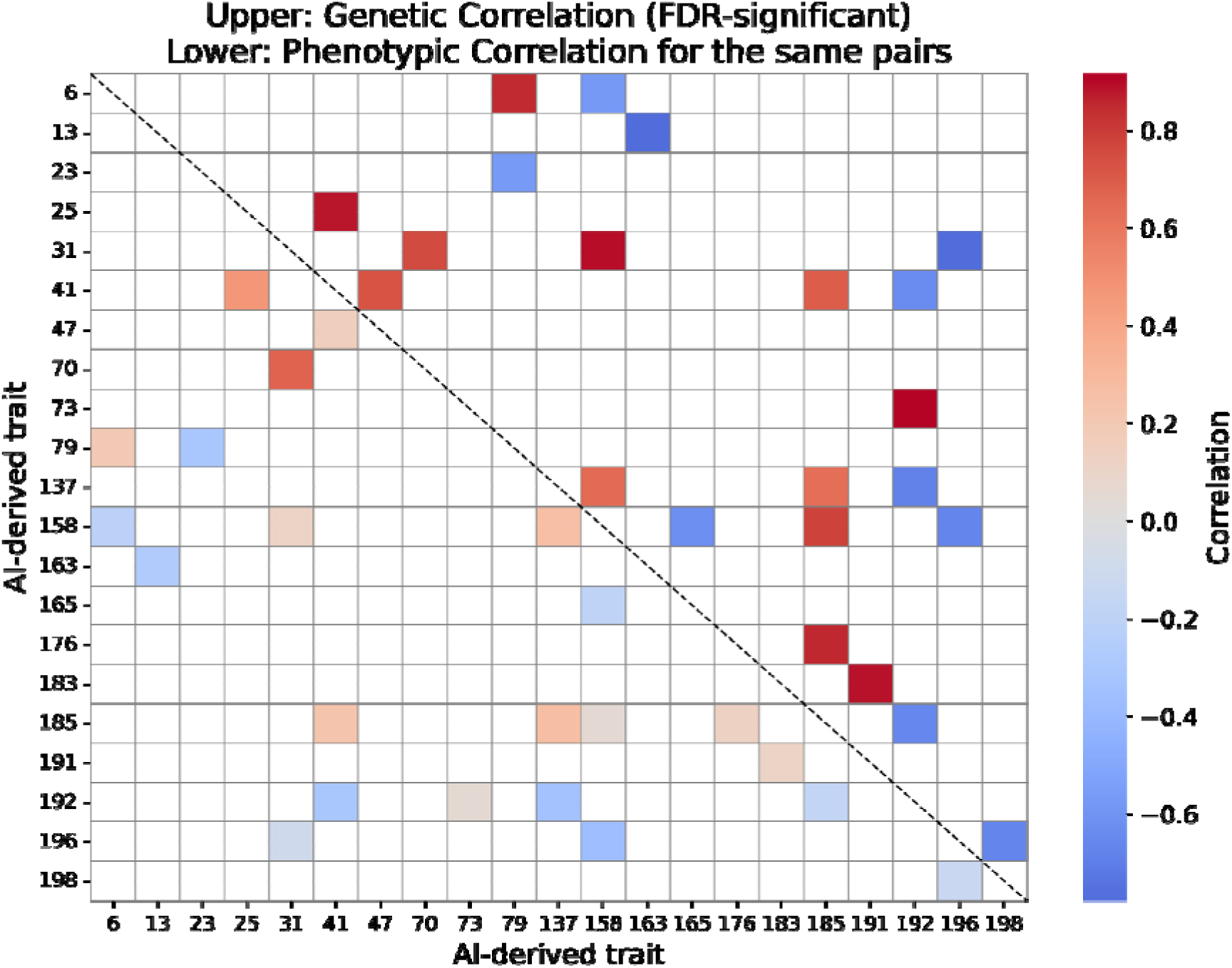
LDSC-based genetic correlation between AI-derived facial traits with FDR-significant associations, and the corresponding phenotypic correlation.

Additionally, we carried out a genetic correlation analysis between the 200 AI-derived facial traits and various disease traits (Figure S7). Nominal significance correlations were seen for diseases such as ADHD, bipolar disorder, major depressive disorder, other Tic disorders and Tourette syndrome, schizophrenia, and alcohol use disorders; however, none of them remained statistically significant after multiple testing correction.

### C-GWAS comparison of AI-derived traits and landmark-based traits

To explore the impact of our novel AI-based versus previously used landmark-based facial phenotyping strategies on C-GWAS outcomes, we compared genetic associations identified with C-GWAS using AI-derived facial traits with those obtained from landmark-distance-based traits previously applied in RS□. This comparison was restricted to 3,761 RS individuals with both, AI-derived and landmark-based facial data available, resulting in a reduced sample size compared to the primary AI-based RS analysis (N = 3,995). The QQ plot indicates that the C-GWAS based on AI-derived facial traits exhibits higher statistical power than that of landmark-based facial traits (Figure 10). In particular, the C-GWAS based on AI-derived facial traits identified 13 significant loci, whereas that from landmark-based traits revealed 10 significant loci, with only three loci overlapping between the two analyses (Figure 10). This limited overlap in the same samples suggests that the two phenotyping approaches capture largely distinct aspects of facial shape variation with different underlying genetic basis.

**Figure 10:**
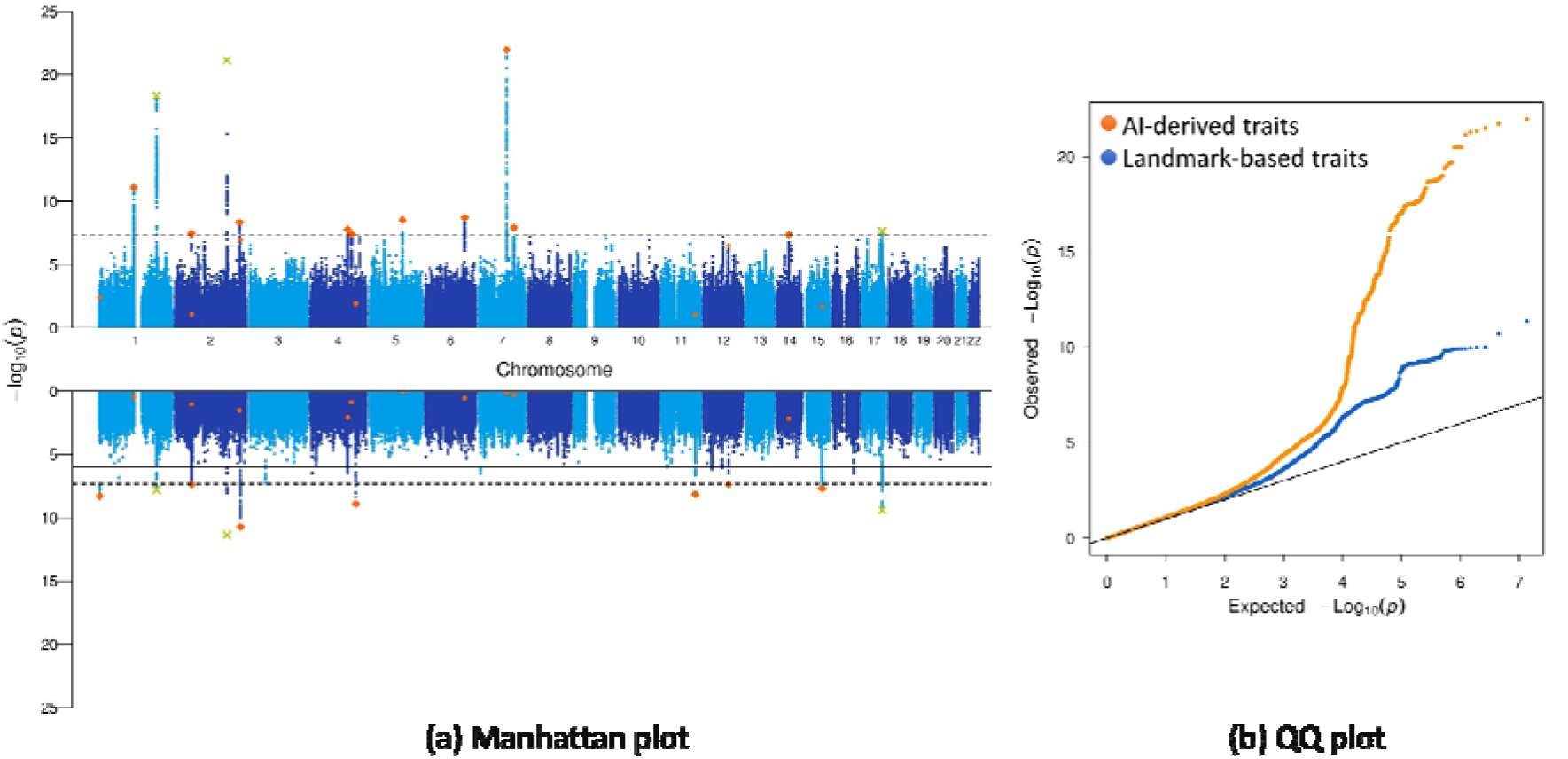
C-GWAS comparison of AI-derived and landmark-based facial traits. a) Manhattan plot: AI-derived (top) vs. landmark-based (bottom) C-GWAS outcomes. Green-crosses denote SNPs that were significant in both analyses. Red-dots denote SNPs that were only significant in one of the two analyses; b) QQ plot: AI-derived (orange) vs. landmark-based (blue) results.

## Discussion

In this study, we developed an innovative, comprehensive, integrated AI-driven computer pipeline to analyse image-based complex traits and to unveil their genetic basis by using multi-cohort data. The proposed pipeline is fully data-driven; hence, no a-priory knowledge is required, whereas other phenotyping strategies, such as landmarking, rely on prior knowledge. Implementing the FL strategy into AI-based phenotyping provides a strong advantage for studying image-based complex traits underlying many genes of small effect size, where inter-institutional multi-cohort studies are required to reach the necessary large sample size, but individual image sharing between institutions often is restricted by stringent privacy regulations.

The effectiveness of our analysis pipeline was demonstrated with the example of 3D facial images. Applying the pipeline to a moderate sample size (N=7,309) from two cohorts, and using 200 AI-derived facial traits, we identified 43 genetic loci, of which 31 represent previously identified genetic face loci. Notably, for known genomic loci, the highlighted patterns in the AI-based facial heatmaps align remarkably well with the effects described for these loci in the literature^5^. For example, *PRDM16*, *PAX3,* and *CRB1* were previously described to be associated with the nose apex^5,37^, nasal bridge^9^ and chin dimples^5^, respectively, which agrees with our XAI-based facial visualization results of the genetic associations we re-discovered for these loci in our data. The 31 re-discovered identified loci also showed associations with various health conditions, such as lung function (*ALX1, CASC17*)^40,41^, asthma (*CRB1*)^42^, coronary artery disease (*CRB1*)^43^, liver enzyme levels (*SEM1*)^55^, insomnia (*CRB1*)^44,45^, Alzheimer’s disease (*EYA4*)^51^, brain morphology (*PAX3, SEM1, ALX1*)^52,53^. Next to providing confirmatory evidence for these previously identified loci, our results demonstrates that our approach using AI-derived facial traits is able to re-identify loci previously discovered by use of substantially different ways of facial phenotyping used previously.

In addition to re-identifying 31 previously known genetic face loci, our approach also highlighted 12 genetic loci that were unknown yet to be involved in facial variation. This finding is remarkable given the moderate sample size we used, which was smaller than in some previous studies. Our high re-discovery rate of 72% provides general confidence that the newly identified loci reflect true positive findings. Moreover, empirical confirmation for 30 of them was provided by our replication analysis in independent European samples. The 12 novel loci are linked to other traits, such as QRS duration (*HEATR5B*)^54^, brain morphology (*HEATR5B, ANK3*)^57^, total cholesterol measurement (*UVRAG*)^62^, and Alzheimer disease (*SCAPER*)^63^. These traits are known to be associated with the human face^64,66,67,68^, and therefore, these findings further justify the robustness of our AI-based approach and validate the identification of these novel loci. Functional studies and experimental validation are needed to explore how these genetic variants influence both facial traits and their associated phenotypic pathways.

### C-GWAS outperformed single-trait GWAS

In the context of AI-based phenotyping followed by GWAS, all previous image-based GWAS studies adopted the traditional single-trait GWAS approach on AI-derived traits from autoencoders, e.g. single-trait GWAS on brain^14^ and cardiac shape^15^. Specifically, one study demonstrated that autoencoder-derived traits from 3D facial meshes have limited performance in single-trait GWAS^80^. In that study, the autoencoder-derived traits did not outperform conventional landmark-distance-based traits in single-trait GWAS, highlighting the limited gain of AI–derived traits under a univariate testing framework. Our results not only confirm this limitation but also provide insight into how it can be addressed using C-GWAS, which leverages multiple AI-derived traits. As shown in Figure 4, in our dataset, single-trait GWAS identified significantly fewer hits, i.e., only one, compared to the many multi-trait GWAS hits (43) achieved with C-GWAS. The underlying reason for this phenomenon partly lies in the unsupervised nature of the applied AI models, which cannot ensure that the AI-derived traits (i.e., basis vectors 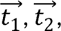 etc. in Figure 3) correspond to facial phenotypes that are strongly associated with a SNP. For example, the nose apex is known to be associated with *PRDM16*^5,37^, but we failed to identify it among any of the 200 traits in our single-trait GWAS. This is because the variability of the nose apex was precisely captured by another vector instead of any of the 200 basis vectors. From this perspective, it is important to search—guided by C-GWAS—for additional facial phenotypes represented by other vectors beyond the 200 basis vectors. Moreover, given that previous face GWASs and C-GWASs based on other ways of facial phenotyping have already discovered hundreds of genetic loci^5,6,9,37,38,46,47,48,81^, of which 31 were re-discovered here, our approach of AI-based phenotyping coupled with C-GWASs allows to unveil new genetic loci involved in facial shapes because they were likely not covered by the previously used facial phenotypes. Thus, for the example of the human face at least, our novel approach reflects a synergistic approach to understand the genetic architecture underlying variation of facial shape more completely.

### FL for alignment of AI-derived traits across cohorts

The alignment of AI-derived traits across multiple cohorts poses a significant challenge in data-driven AI-based phenotyping due to variations in data distributions. Traditionally, this challenge has been addressed through centralized training strategies, where data from multiple cohorts are shared and combined into a single source for model training. However, this approach is not in line with the practise of GWAS meta-analysis in multi-cohort studies needed to achieve larger sample size, where researchers typically prefer to share GWAS summary statistics data between collaborating institutions instead of individual data. Moreover, sharing individual-level images, especially those that allow individualization such as facial images, often have strong hurdles in sharing across institutions and countries, which sometimes cannot be solved and thus such image data cannot be shared. To overcome this limitation, we integrated an alternative training strategy: federated learning (FL) in our analysis pipeline. Our proposed FL strategy has demonstrated comparable performance to traditional CT in deriving aligned traits across different cohorts, without the need for image sharing between different sites. This property of our pipeline facilitates collaborative C-GWAS efforts involving multi-cohort data from different institutions.

Data harmonization across cohorts is a key consideration in multi-cohort genetic studies integrating imaging-derived phenotypes. In the present application, potential site-related effects were minimized at both the imaging and genetic levels. All facial images were acquired using the same 3dMD system under standardized protocols, limiting variability due to acquisition hardware or settings, and were subsequently processed using a standardized preprocessing pipeline. For genotype data, all cohorts were processed using a consistent quality-control pipeline and imputed to the same reference panel, i.e., 1000 Genomes^3^, which is standard practice in GWAS and GWAS meta-analyses.

Differences in demographic composition across cohorts (e.g., age or genetic ancestry) may still act as potential confounders. In the present study, no explicit harmonization was applied at the phenotyping stage. Instead, we assessed their practical impact by comparing CT and FL, and the consistency between CT and FL suggests that residual cohort effects did not materially influence phenotype derivation in this setting. More generally, in scenarios where stronger cross-site heterogeneity is present, additional harmonization strategies can be integrated within the federated learning framework, such as cohort-aware normalization or domain adaptation approaches^56^. Prior work has demonstrated the effectiveness of federated learning across heterogeneous sites for other imaging modalities, including 3D volumetric modalities such as brain MRI and computed tomography^56^, as well as 2D RGB modalities such as retinal imaging^79^, supporting its applicability in broader multi-cohort imaging genetics studies.

### Training stability and robustness of AI-derived traits

Autoencoder-based latent representations are known to exhibit variability across training runs due to random initialization and stochastic optimization, resulting in latent axes (i.e., AI-derived traits) that may differ across independent trainings. In our previous work^20^, we explicitly evaluated the robustness of the learned representations by performing multiple independent trainings of the autoencoder and repeating downstream association analyses. Across these independent runs, highly consistent statistical associations and facial heatmap visualizations were observed after combining effects of multiple latent axes, indicating stable and reproducible phenotype-related signals. Although individual latent axes may shift between trainings, the underlying vector directions (e.g., 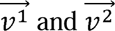 as described in Figure 3) capturing meaningful facial shape variation remain present and can be reliably identified when integrating multiple traits. From this perspective, single-trait GWAS performed on individual AI-derived traits may be more sensitive to training variability, whereas C-GWAS, which aggregates information across multiple AI-derived traits, is expected to provide more robust and reproducible association results. These findings support the robustness of the proposed framework despite inherent training stochasticity.

### Comparison between children and elderly cohorts

The efficacy of our FL strategy was demonstrated through a multi-cohort study of an elderly cohort (mean age=69.6 years, SD=9.5 years) and a children cohort (mean age=10.4 years, SD=1.5 years). Despite the significant age difference between the participants of these two cohorts, the combination of data in the joined C-GWAS led to an improvement in power for most of the identified genetic loci. For instance, *PRDM16*, *CRB1*, *MTX2*, and *EYA4* exhibited similar patterns in facial heatmaps across the two cohorts, and their statistical power was enhanced after the meta-analysis (Figure 6). This highlights the success of our proposed pipeline in conducting GWAS meta-analysis across multiple cohorts.

However, we also observed a decrease in power for five genetic loci after the meta-analysis. For example, rs2296926 (*TRMT5,* p-value: 4.1E-08 in RS), rs111520336 (*BRINP1*, p-value: 5.6e-10 in GenR) and rs36109172 (*GHR*, p-value: 4.1E-08 in GenR) showed different facial patterns between the two cohorts (Figure 7). While they were genome-wide significant in either the elderly cohort or the children cohort alone with separate C-GWASs, they did not retain statistical significance after meta-analysis in the combined C-GWAS. We hypothesize that this discrepancy may be attributed to the large difference in age between the two cohorts. This hypothesis is substantiated by some of the identified loci. For instance, *BRINP1* is strongly associated with body height, adolescent idiopathic scoliosis, and cognitive function^69,70,71^, while *TRMT5* is highly related to bone density^72^, and *GHR* is closely linked to body height and growth hormone receptor measurement^69,74^. These traits, like facial morphology, are age-dependent and likely undergo dynamic changes across the lifespan. Age-related differences may alter the expression or relevance of genetic loci, potentially shifting their impact on facial phenotypes at different stages of life.

### Strengths and limitations

Different from other existing multi-trait GWAS pipelines^5^, our pipeline that finally uses C-GWAS, retains conventional single-trait GWAS summary statistics, thus offering the opportunity to perform genetic correlation analysis with other phenotypes. Our AI-derived facial traits demonstrate genetic correlations with various diseases; however, the obtained correlation is weak and lacks statistical significance. This result is in line with prior findings about the weak genetic correlation between diseases and normal facial traits^64^. Interestingly, we identified high genetic correlation among the 200 AI-derived facial traits, suggesting the pleiotropic effects of a SNP on multiple traits. Therefore, applying C-GWAS to combine multiple AI-derived traits is crucial to improve the power by aggregating these shared genetic signals, especially when sample size is modest. As shown in Figure 4, multi-trait GWAS (43 hits) facilitated through the C-GWAS approach, yielded significantly more hits than the single-trait GWAS (one hit).

A notable limitation of this study lies in the significant disparity in age distribution between the two cohorts, which likely restricted the extent of power enhancement through meta-analysis. Nevertheless, we anticipate that the performance of our pipeline would be notably improved in scenarios where the age distribution across cohorts exhibits greater similarity. Based on the compelling evidence from this study, we anticipate that our novel analysis pipeline will emerge as a crucial tool for facilitating inter-institutional collaborations involving multiple cohorts to unveil the genetic basis of image-based complex phenotypes, while effectively addressing data privacy concerns by preventing inter-institutional sharing of individual images.

## Methods and materials

### Study population

#### Rotterdam Study

The Rotterdam Study (RS) is a population-based cohort study of 14,926 participants aged 45 years and older, living in the same suburb of Rotterdam in the Netherlands^1,17^. The present study includes 3,995 participants (mean age=69.6 years, SD=9.5 years) of Dutch European ancestry, for whom high-resolution 3-dimensional digital face photographs were taken. The raw 3D facial images were acquired using a 3D photographic scanning system manufactured by 3dMD (Atlanta (GA), USA). Participants were asked to keep their mouths closed and adopt a neutral expression during the acquisition of the 3D scans.

Microarray-based genotyping of genome-wide single nucleotide polymorphisms (SNPs) was carried out according to the manufacturer’s instructions with the Infinium II HumanHap 550K Genotyping BeadChip version 3 (Illumina, San Diego, California USA) at the Human Genomics Facility (HuGe-F) of the Department of Internal Medicine of Erasmus MC (www.glimdna.org)^2^. All SNPs were imputed using MACH software (www.sph.umich.edu/csg/abecasis/MaCH/) based on the 1000-Genomes Project reference population information^3^. After all quality controls, the current study included a total of 6,728,663 autosomal SNPs (MAF > 0.01, imputation R2 > 0.8, SNP call rate > 0.97, HWE > 0.0001).

The Rotterdam Study has been approved by the Erasmus MC Medical Ethics Committee (registration number MEC 02.1015) and by the Dutch Ministry of Health, Welfare and Sport (Population Screening Act WBO, license number 1071272–159521 PG). The Rotterdam Study has been entered into the Netherlands National Trial Register (NTR; www.trialregister.nl) and into the WHO International Clinical Trials Registry Platform (ICTRP; www.who.int/ictrp/network/primary/en/) and under shared catalogue number NTR6831. All participants provided written informed consent to participate in the study and to have their information obtained from treating physicians.

#### Generation R Study

The Generation R Study (GenR) is an ongoing population-based cohort study of pregnant women and their children from fetal life onwards^18^. The goal is to identify early environmental and genetic causes leading to normal and abnormal growth, development, and health. All women living in the study area of Rotterdam in the Netherlands, who gave birth between April 2002 and January 2006 were eligible. The present study includes 3,314 participants (mean age=10.4 years, SD=1.5 years), for whom high-resolution 3-dimensional face digital photographs were taken using the 3dMD system. The participants were of multi-ethnic ancestry^35,36^, and the largest groups were Dutch-Europeans (N=2,026, 61.1%), non-Dutch-Europeans (N=243, 7.3%), Turkish (N=186, 5.6%) and Moroccans (N=175, 5.3%).

Microarray-based genotyping^19^ was performed at the Human Genomics Facility (HuGe-F) of Erasmus MC. All SNPs were imputed using MACH (version 1.0.15) software based on the 1000-Genomes Project reference population information^3^. After all quality controls, the current study included a total of 7,416,225 autosomal SNPs (MAF > 0.01, imputation R2 > 0.8, SNP call rate > 0.98, HWE > 1.10e–6).

The Generation R Study was approved by the Erasmus MC Medical Ethics Committee (MEC) (MEC 198.782/2001/31), and written informed consent was obtained from all participants, either themselves or from their guardians/parents.

### Facial traits from the AI-based modelling

The raw 3D facial image data were built into a template-based data set following a registration pipeline^20^. A 3D mesh autoencoder, whose architecture and implementation were previously introduced and validated in our earlier work^20,21^, was applied to learn compact facial representations in a self-supervised manner for dimensionality reduction. More specifically, the autoencoder consists of a four-layer encoder and a four-layer decoder that perform non-linear feature mapping using spiral convolution operations followed by exponential linear unit (ELU) activations^21^. The encoder compresses high-dimensional facial meshes into low-dimensional latent representations, while the decoder reconstructs the original facial geometry from these representations. The network was trained by minimizing the reconstruction error between the input and reconstructed facial images, such that the latent representations capture the major modes of facial morphological variation. Based on empirical evaluation reported in our previous study²□, the dimensionality of the latent space was set to 200 as a trade-off between reconstruction accuracy and model complexity^20^. In this phenotyping step, the autoencoder was trained using the full dataset in an unsupervised manner. We defined these 200 learned representations as AI-derived facial traits, which were used in the GWAS as phenotypes.

### Alignment of AI-derived traits across multiple cohorts via federated learning

Federated learning (FL)^65^ (also known as collaborative learning) is a machine learning technique that trains a model via multiple independent sessions, each using its own dataset. This approach stands in contrast to traditional centralized training (CT) techniques, where local datasets are merged into one training session^22^. A server was established upon clients of multiple (M) institutes, with a randomly initialized global AI model to be trained. In each round of the training, the global model is updated via following steps: 1) The server sends the global model to client #1, and then client #1 updates the received model with its own local data; 2) Client #1 sends the updated model back to the server; 3) repeat 1) and 2) for the rest (M-1) clients; 4) repeat 1), 2) and 3) for a prefixed number of rounds of the training. After the training is finished, a final global model was sent to clients. With these settings, we are able to train the proposed AI model simultaneous among multiple cohorts without sharing individual-level image data. Most importantly, since the AI model is jointly optimised with facial image data from multiple institutes, the representing facial morphology of each trait is aligned across institutes (Figure 1). This allows meta-analysis on traditional GWAS summary statistics of cohort data across institutes as well as C-GWAS.

To better understand the efficacy of the FL training strategy, we compared the performance of the FL and CT approaches in terms of the reconstruction error (Equation 1) between input and reconstructed facial meshes. In this comparison, the FL and CT models used the same random data split: 80% for training and 20% as an independent test set, with performance evaluated on the same test data. The reconstruction error was calculated as follows:

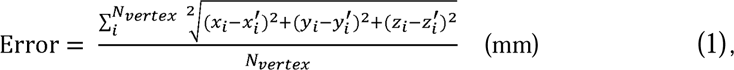

where *N_vertex_* is the number of vertices of the facial mesh, (*x_i_*, *y_i_*, *z_i_*) is a vertex of the input facial mesh while (*x_i_′*, *y_i_′*, *z_i_′*) is the corresponding vertex in the reconstructed facial mesh.

### Single-trait GWAS, meta-analyses, and C-GWAS

In RS, traditional single-trait GWASs were conducted separately for each and every of the 200 AI-derived traits using a total of 6,728,663 SNPs. GWAS was performed using linear regression under the additive genetic model while adjusting for sex, age, and genetic ancestry derived from the first four genomic PCs using Plink^4^. In GenR, GWASs were conducted separately for each and every of the 200 AI-derived traits using a total of 7,416,225 SNPs using HASE^23^ while adjusting for sex, age, and the first twenty genomic PCs. More genomic PCs were considered for GenR compared to RS because of the multi-ethnic nature of the GenR cohort relative to the Dutch-European RS cohort. Inverse variance fixed-effect meta-analysis were carried out to combine GWAS results of RS and GenR for each of the 200 AI-derived traits, which included 7,733,921 autosomal SNPs with physical positions aligned according to 1NCBI human genome sequence build GRCh37.

Using the summary statistics from the single-trait GWASs of the 200 AI-derived facial traits, we performed C-GWAS^6^. The C-GWAS R library is freely available at https://github.com/Fun-Gene/CGWAS. Details of the C-GWAS method have been described elswhere^6^. In brief, the null hypothesis (H0) for a tested SNP is the absence of any allelic effect on all traits, and the alternative hypothesis (H1) is that its allelic effects deviate from 0 for at least one of the multiple traits tested. C-GWAS incorporates two different tests that originated from either the effect-based inversed covariance weighting, or the truncated Wald test to maximise statistical power. All resultant P-values from C-GWAS are adjusted using the getCoef function implemented in C-GWAS, which performs simulations (n simulations = 1e8) to guarantee that the null distribution of C-GWAS follows the uniform distribution in all quantiles. A common approach for handling correlated traits in GWAS involves performing separate GWASs for each single trait and selecting the minimum p-value (unadjusted MinGWAS). However, unadjusted MinGWAS is inflated under the null hypothesis (no effect on any trait), requiring stricter significance thresholds or adjustments for multiple correlated traits. In this study, we adjusted our unadjusted MinGWAS, i.e., minimal values of meta-analysis of 200 AI-derived traits GWASs, using the getCoef function in C-GWAS. This function employs simulation analyses to ensure that both C-GWAS and the adjusted MinGWAS follow a uniform distribution under the null. Consequently, our MinGWAS refers to the getCoef-calibrated minimum p-value, allowing direct comparison between the outcomes of C-GWAS, MinGWAS, and any single-trait GWAS. Thus, traditional genome-wide significance threshold of 5e-8 corresponds to our study-wide significance threshold for both C-GWAS and MinGWAS. Here, we used the default parameters of C-GWAS and applied C-GWAS to the summary statistics of the 200 AI-derived traits GWASs in the meta-analysis of GenR and RS. Moreover, although not part of the core process of our proposed pipeline, we conducted additional C-GWAS separately in GenR and RS, to further illustrate how meta-analysis with our pipeline can improve the power.

### Replication of C-GWAS outcomes

Aiming to replicate the C-GWAS derived top SNPs, we reached out and obtained the GWAS summary statistics (GWAS Catalog: GCP000044)^5^ from 8,246 individuals of European descent from a combination of three independently collected datasets from the United States (US; N = 4,680) and the United Kingdom (UK; N= 3,566). The data contains 126 sets of GWAS summary statistics corresponding to the two sets of 63 facial segments of US-driven or UK-driven meta-analyses respectively. These facial segments are derived as linear combinations of facial vectors within distinct regions, and they do not directly correspond to our AI-derived traits based on whole facial vectors. Due to the differences in phenotyping methods between the discovery and replication studies, we used minimal p-values and C-GWAS embedded function getCoef to combine the evidence of replication from 126 GWASs and account for the correlation between facial segments. In brief, we first integrate 126 GWASs to one GWAS (MinGWAS) by taking the minimal p-values for each SNP. This is to test the null hypothesis that the SNP effect on all 126 facial segments are 0. Then we calibrate MinGWAS using getCoef function in C-GWAS, which guarantees the null of calibrated MinGWAS is the same as uniform 0-1 distribution and any of conventional GWAS^6^. We selected calibrated MinGWAS p-values of 43 lead SNPs as replicated evidence. p<0.05 indicates that the replicated SNP has effect on at least one facial segment with study-wide nominal significance.

### Comparison of C-GWAS from AI-derived and landmark-based facial traits

To directly compare AI-based and landmark-based facial phenotyping approaches in their genetic outcomes, we conducted single-trait GWAS separately for AI-derived traits and landmark-derived distance traits using the subset of RS participants for whom both phenotyping modalities were available (N = 3,761). The AI-derived traits were identical to those used in the primary analyses, and the AI model was not retrained for this comparison. The landmark-based distance phenotypes consist of 78 traits derived from 13 facial landmarks, as detailed in our previous study^9^. After conducting single-trait GWAS, we subsequently applied C-GWAS to the summary statistics of the 200 AI-derived traits, and the 78 landmark-based traits and compared the outcomes.

### Visualizing genetic association effects on the human face

Explainable artificial intelligence (XAI) aims to enhance the interpretability of machine learning models and their learned representations, particularly in biomedical image analysis^50^. In imaging genetics, such interpretability can facilitate biological insight by linking learned representations to anatomically meaningful structures. While XAI is commonly associated with explaining predictions in supervised learning tasks, the present work operates in a different setting—unsupervised representation learning for gene discovery and feature localization.

Here, explainability is achieved at the level of the learned latent trait space. The autoencoder provides a structured vector space in which different vector directions represent distinct modes of facial morphological variation, which may reflect global morphological structure, localized deformation, or variation associated with specific biological factors (e.g., genetic variants, age or sex)^39,73^. By estimating SNP-specific effect sizes across latent traits, we constructed SNP-guided direction vectors that represent the composite phenotypic effect of each genetic variant. Traversing the latent space along these directions enables controlled decoder-based reconstruction, thereby isolating and visualizing the anatomically localized facial features associated with an identified SNP. In this sense, our XAI framework introduces vector-wise insights into the latent space and provides representation-mediated biological explainability of genetic association signals.

Specifically, we performed a per-SNP facial visualization for the C-GWAS-derived top SNPs. After the training of the autoencoder, each input facial mesh is represented by a latent datapoint (blue points in Figure 3) in the AI-derived trait space. The trait space can be viewed as an n-dimensional vector space □^n^ spanned by basis vectors 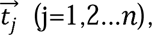 which form a canonical (not necessarily orthogonal) basis. Any vector in this space can be expressed by 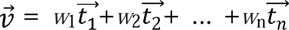 as a linear combination of basis vectors using their weightings *w_j_* (j=1,2…n)^39,73^ (Figure 3). By decoding a selected trait (e.g., t_1_) while keeping the rest unchanged, the decoder can perform sampling along an axis (e.g., 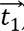) in the trait space, and to reconstruct a series of facial shapes showing deformation related to the change of the selected trait (Figure 3). Particularly, by selectively decoding a subset of traits with weightings *w*_1_, *w*_2_, …, *w*_n_, the decoder can perform sampling along vector 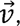 and to reconstruct a series of facial shapes showing facial deformation related to the linear combination of multiple traits. A facial heatmap is calculated to better visualize the facial deformation, where outward deformation, with respect to the geometric center of the head, were in red color. C-GWAS combines multiple traits based on single-trait summary statistics of all generated facial traits. Therefore, given 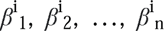 as betas of n traits for SNP in the single-trait GWAS, by decoding and sampling latent datapoints along 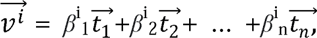 we are able to visualise facial phenotypes associated with SNP^i^ in the C-GWAS results. The range of the sampling is mean ± SD of latent datapoints from the study population. Notably, we only selected nominal-significant (p<0.05) traits for this visualization.

### Heritability and genetic correlation analyses

We performed LDSC^7^ on the meta-analysis GWASs to estimate the genome-wide heritability for each of the 200 AI-derived traits. After the estimation of heritability on facial traits, we mapped the heritability of the traits to the human face. Implementation details are described in Supplementary Figure S4 and Figure S5. We applied LDSC to compute the genetic correlation between the 200 AI-derived traits. Furthermore, we estimated genetic correlation (rg) between face shape and brain diseases on a summary level, by using publicly available GWAS summary statistics for 11 diseases and disorders including anxiety disorder^24^, attention deficit hyperactivity disorder (ADHD)^25^, autism spectrum disorder^26^, Alzheimer’s disease^27^, bipolar disorder^28^, cross disorder^29^, eating disorder^30^, major depressive disorder^31^, other Tic Disorders and Tourette syndrome^32^, schizophrenia^33^, and alcohol use disorders^34^. We applied LDSC^8^ to calculate rg for each pair of face and disease GWAS.

## Acknowledgement

We thank all cohort participants or their legal representatives for their participation.

## Author contribution

X.L. and Z.X. contributed equally to this work and performed the main analyses, with X.L. leading AI phenotyping and federated learning and Z.X. leading GWAS, meta-analysis, and C-GWAS. F.L., T.N., and E.B.W. contributed to data resources, study design, and interpretation. M.K. and G.V.R. conceived and supervised the study and contributed to interpretation. X.L. and Z.X. drafted the manuscript. All authors reviewed and approved the manuscript. M.K. and G.V.R. jointly supervised this work.

## Code availability

The scripts for the proposed methodological pipeline ‘flaipGWAS’, along with a demo of the 3D face mesh application, are available on GitLab https://gitlab.com/radiology/compopbio/fl_based_ai_phenotyping_for_gwas. C-GWAS is implemented as an open-source R package available at https://github.com/Fun-Gene/CGWAS. The HASE is implemented as an-open-source python package available at https://github.com/roshchupkin/HASE. LDSC is implemented as an open-source command line tool available at https://github.com/bulik/ldsc. The mesh processing library is implemented as an open-source python package available at https://github.com/MPI-IS/mesh. The source code of 3D mesh autoencoder is available at https://github.com/sw-gong/spiralnet_plus and the implementation scripts for AI-based facial phenotyping are provided in our flaipGWAS package. The source code of the NVFlare framework for federated learning is available at https://github.com/NVIDIA/NVFlare and the implementation scripts for AI-based facial phenotyping are provided in our flaipGWAS package. Other software used in data analysis includes: Plink (https://www.cog-genomics.org/plink2/), FastGWA (https://yanglab.westlake.edu.cn/software/gcta/#fastGWA), FUMA (https://fuma.ctglab.nl/), and GCTA-Fst (https://yanglab.westlake.edu.cn/software/gcta/#Fst).

## Data availability

The synthetic face mesh data are available at: https://gitlab.com/radiology/compopbio/fl_based_ai_phenotyping_for_gwas/-/tree/main/synthetic_data_demo?ref_type=heads

Summary statistics data of our C-GWAS and MinGWAS are at available at https://zenodo.org/uploads/15783705?token=eyJhbGciOiJIUzUxMiJ9.eyJpZCI6Ijk1OTdiNTk3LTI5Y2QtNGFkZS05NGY4LThkYTFmYTg5Y2M0ZCIsImRhdGEiOnt9LCJyYW5kb20iOiIzNDFhOTBmMWNhNGE5ZmI4MmEyNDg2MjBmZTEyNmNjMiJ9.yj_JRC6Tjzeh3P_Yp-WELAjLrEpJL0uD8JfnUn8fvYAK2GfTzM66bUGe5MZQ8f6SBNgBlOFSe_0nDWpVPHmasQ.

Single-trait GWAS summary statistics of our RS and GenR cohorts, before and after meta-analysis, are available at https://zenodo.org/records/15786879?preview=1&token=eyJhbGciOiJIUzUxMiJ9.eyJpZCI6ImQ5YzU4Mzk0LTRhY2UtNDEwNi1hNzk2LTQzOGUzOWNlYWZhNiIsImRhdGEiOnt9LCJyYW5kb20iOiI2ODI1ODRlYzAwNTAzZDQzYzVjMTI4YmRiMDNmNDEwNyJ9.FNRBRsodum7AF5ll9D6TDcBuGl_BQnxa0QPAlypDU4gXdD3cfYkVSsQcjZe_hk2OZEmZ_B_GpBKrPKcW4QtG-w

GWAS summary statistics used for the replication analysis are publicly available at GWAS Catalog (GCP000044). GWAS summary statistics for 12 diseases and disorders are publicly available, including anxiety disorder^24^ (10.6084/m9.figshare.16602218), ADHD (10.6084/m9.figshare.19383299), autism spectrum disorder^26^, (10.6084/m9.figshare.14671989) Alzheimer’s disease^27^ (https://vu.data.surfsara.nl/index.php/s/l7aiRr1UEgdoJfZ) bipolar disorder^28^ (10.6084/m9.figshare.14102594), cross disorder^29^(10.6084/m9.figshare.14672034), eating disorder^30^ (10.6084/m9.figshare.14671980), major depressive disorder^31^(10.6084/m9.figshare.14672085), other Tic Disorders and Tourette syndrome^32^ (10.6084/m9.figshare.14672232), schizophrenia^33^ (10.6084/m9.figshare.14672154), and alcohol use disorders^34^ (10.6084/m9.figshare.14672193).

Access to individual-level data from the GenR and RS cohorts used in this study is restricted to researchers in direct collaboration with the cohort owners. For RS, collaboration requests can be directed to Frank J.A. van Rooij at Erasmus MC; For GenR collaboration requests can be directed to the data manager at Erasmus MC. Responses to collaboration requests will be answered in a timely fashion, typically one calendar month.

## Notes

**Funding sources:** The Rotterdam Study is funded through unrestricted research grants from Erasmus MC University Medical Center Rotterdam, Erasmus University Rotterdam, the Netherlands Organization for the Health Research and Development (ZonMw), the Research Institute for Diseases in the Elderly (RIDE), the Ministry of Education, Culture and Science, the Ministry for Health, Welfare and Sports, the European Commission (DG XII), and the Municipality of Rotterdam. The Generation R Study is conducted by the Erasmus MC in close collaboration with the School of Law and Faculty of Social Sciences of the Erasmus University Rotterdam, the Municipal Health Service Rotterdam area, Rotterdam, the Rotterdam Homecare Foundation, Rotterdam, and the Stichting Trombosedienst & Artsenlaboratorium Rijnmond (STAR-MDC), Rotterdam. Gennady V. Roshchupkin supported by the ZonMw Veni grant (Veni 1936320).

**Conflicts of Interest:** The authors declare not to have a conflict of interest with this work.

### Competing Interest Statement

The authors have declared no competing interest.

### Summary of Updates

Revision based on reviewers comments, and adapted to PNAS format

