## Supplementary for "Genetic Insights of Image-Based Traits: Analysis Pipeline for AI-based Phenotyping, Combined-GWAS, and Federated Learning with Application to the Human Face"

Supplementary material

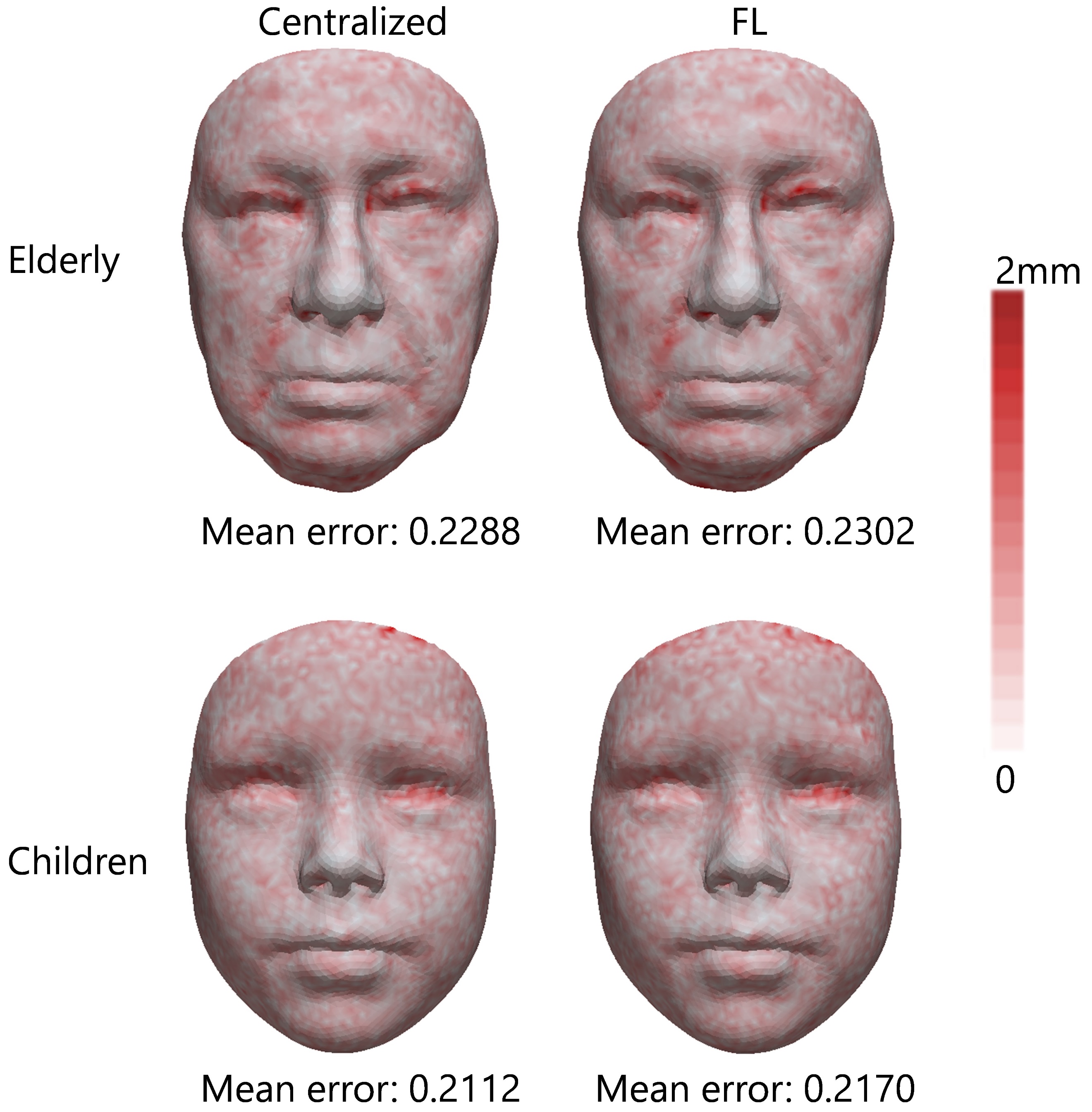

Figure S1: Examples showing reconstruction error of centralized training strategy and federated learning (FL) strategy, for an individual from the elderly cohort and an individual from the children cohort. Numbers below facial images are the mean error of the individual.

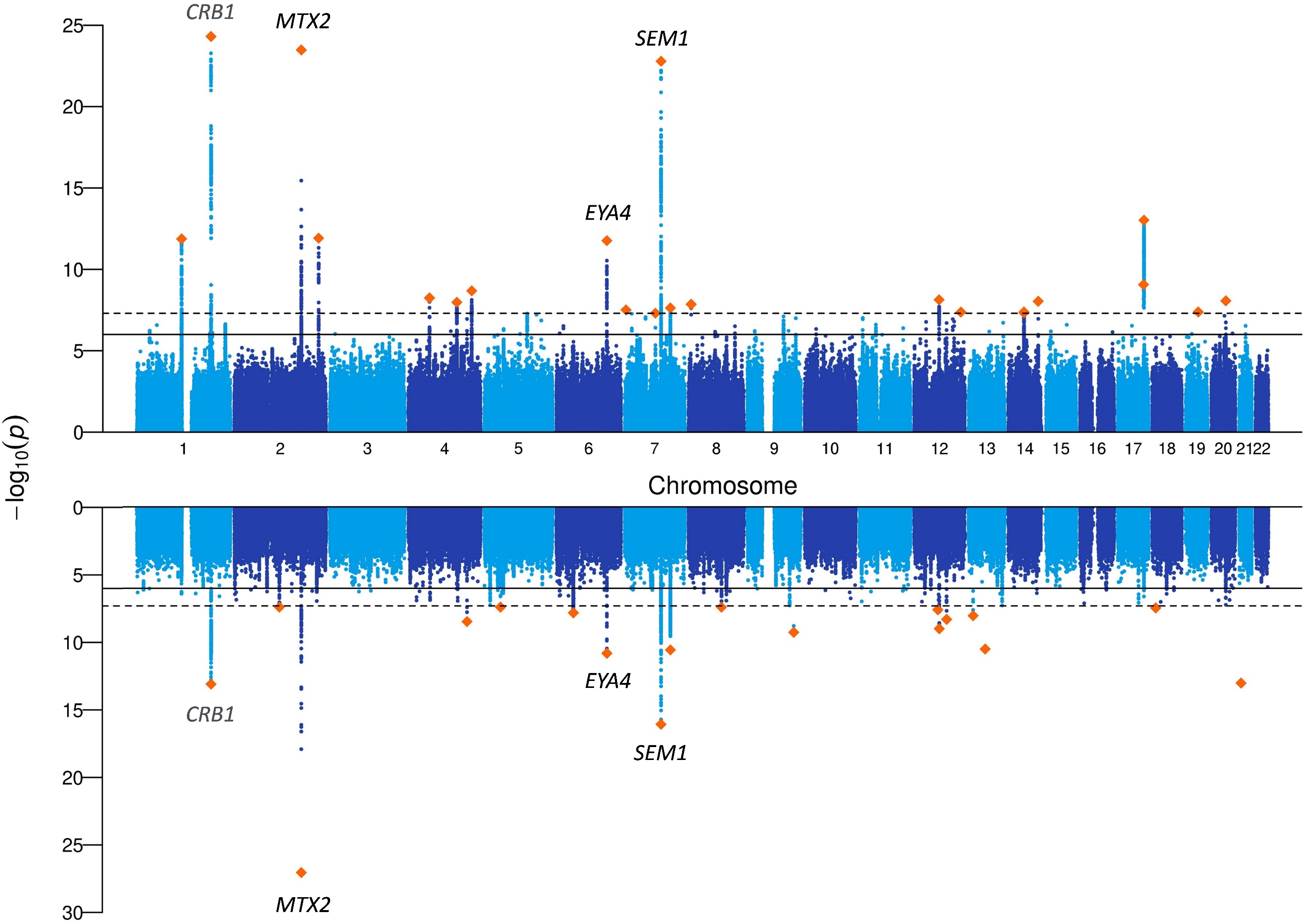

Figure S2: Manhattan plot of the separate C-GWAS for the elderly RS cohort (upper part) and the children GenR cohort (lower part). Four loci i.e., *CRB1*, *MTX2*, *EYA4* and *SEM1* were identified in both analyses.

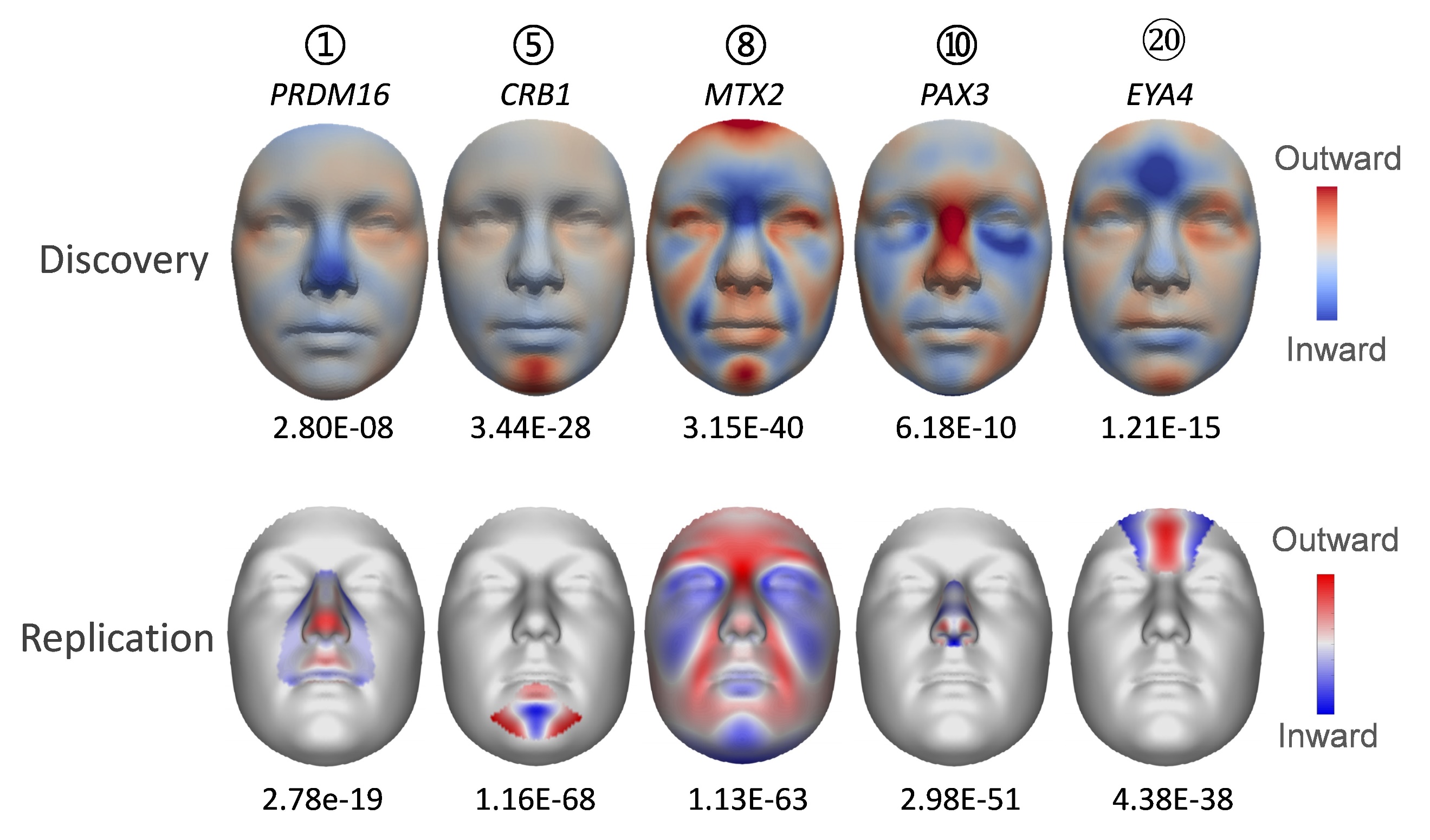

Figure S3: Comparison of facial heatmaps of 5 genome-wide significant genetic loci in the discovery C-GWAS and the replication analysis.

**Implementation details of facial heritability heatmap**

This procedure does not estimate vertex-wise heritability in a statistical genetics sense. Instead, it constructs a facial heatmap that attributes trait-level heritability to facial regions based on explainable-AI–derived spatial representations. The resulting map is intended as an interpretable visualization of where heritable facial variation manifests, rather than a formal heritability estimator at the vertex level.

Below are steps to combine multiple FDR-significant traits to calculate a facial heritability heatmap:

1) Based on the XAI technique, we obtained for each trait a facial relevance map that quantifies the contribution magnitude of each facial vertex to the AI-derived trait (Figure 3). To ensure comparability across traits, each relevance map was transformed to its absolute value and min–max normalized to the range [0,1], yielding a normalized representation map $F_{i}$ for each FDR-significant trait i = 1, 2, ..., n.

2) Add heritability weighting onto each normalized representation map:

$H_{i}=h_{i}F_{i}$, i = 1, 2, …, n

where $h_{i}$ is the heritability of each AI-derived trait.

Thus, $H_{i}$ is the heritability map of each AI-derived trait.

This step assumes that the global SNP-based heritability of a trait can be spatially attributed according to the relative importance pattern captured by the XAI relevance map.

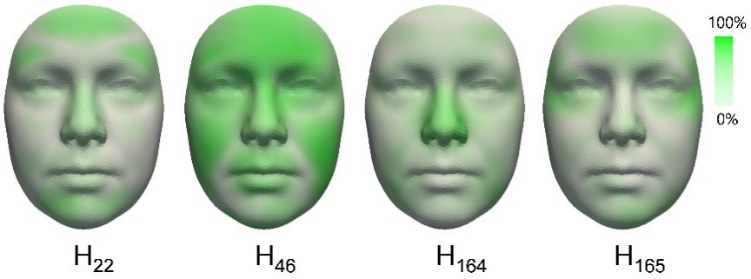

Figure S4: Heritability map of each AI-derived trait.

3) Convert AI-derived traits into independent components.

Because AI-derived traits are partially correlated, directly aggregating their heritability maps would lead to redundant contributions. We therefore applied independent component analysis (ICA) to decompose the AI-derived traits into statistically independent components, enabling separation of shared and distinct heritable facial patterns.

The n AI-derived traits are thus transformed to independent components $I_{1}, I_{2}, \ldots, I_{m}$ (m components).

4) Project each AI-derived trait onto m components.

The Pearson correlation coefficients between AI-derived traits and independent components are calculated. These coefficients are equivalent to the cosine value between them in the perspective of vector space, i.e., $Corr\left( t_{i},I_{1} \right)= cos<\vec{t_{i}}, \vec{I_{1}}>where Corr(.,.) refers to Pearson correlation.$Therefore, we are able to project the heritability map ($H_{i}$) of each AI-derived trait onto m independent components, and calculate the residuals. The residual term captures the component of the trait-specific heritability map that is not explained by the independent components.

$H_{i}^{p1}$ = $H_{i}cos<\vec{t_{i}}, \vec{I_{1}}>$

$H_{i}^{p2}$ = $H_{i}cos<\vec{t_{i}}, \vec{I_{2}}>$

..

$H_{i}^{pm}$ = $H_{i}cos<\vec{t_{i}}, \vec{I_{m}}>$

$H_{i}^{residual}$ = $H_{i} (1- \sqrt[2]{{cos<\vec{t_{i}}, \vec{I_{1}}>}^{2}+{cos<\vec{t_{i}}, \vec{I_{2}}>}^{2}+\ldots+{cos<\vec{t_{i}}, \vec{I_{m}}>}^{2}}$ )

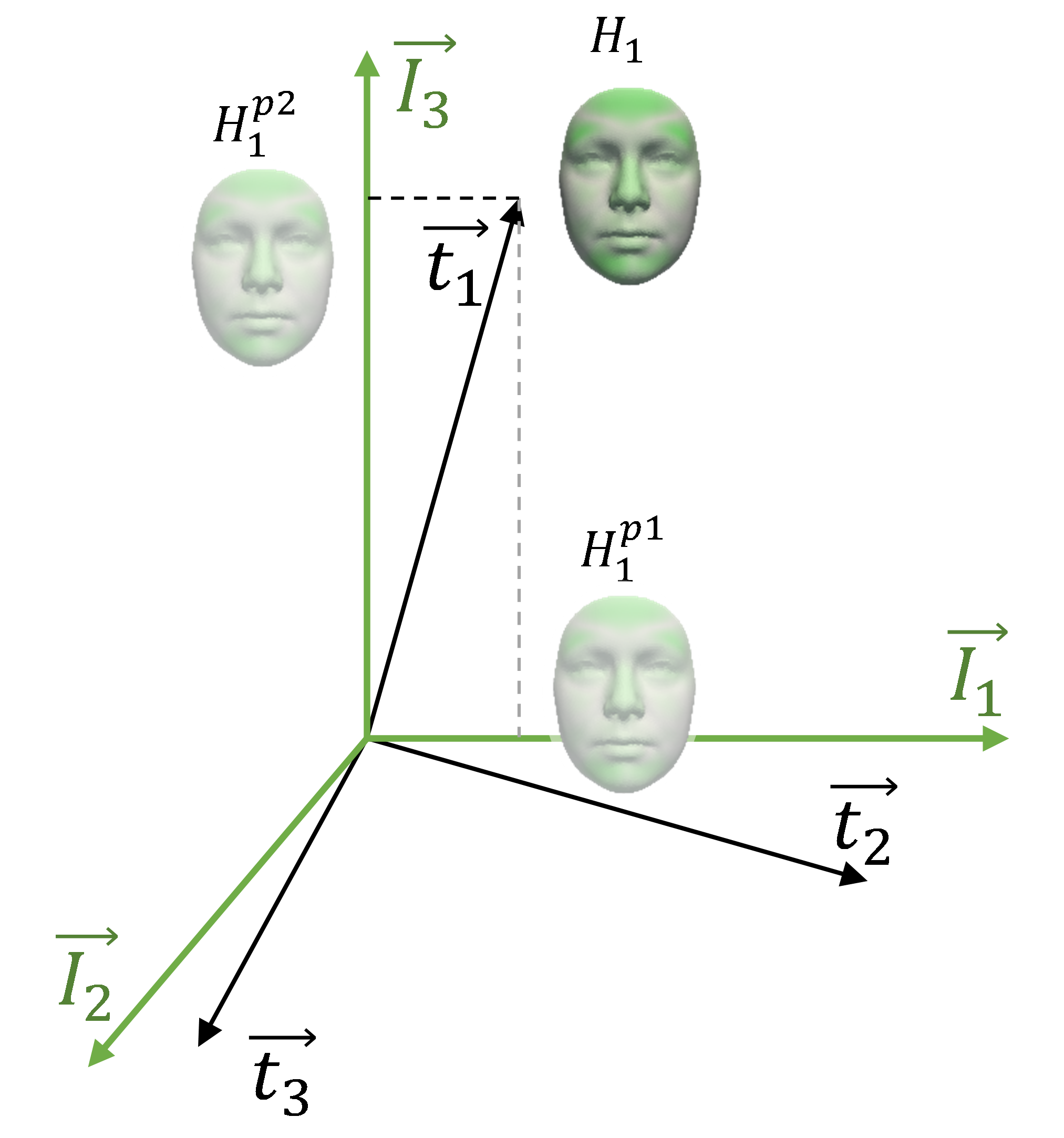

Figure S5: Projecting the heritability map of each AI-derived trait onto m

Components $I_{1}, I_{2}, \ldots, I_{m}$. This figure showcases the example

of projecting $H_{1}$ onto $I_{1}$ and $I_{2}$.

5) Combine heritability of multiple AI-derived traits within an independent component:

$H_{multiTrait}^{p1}=max(H_{1}^{p1}, H_{2}^{p1}, \ldots, H_{n}^{p1}$)

$H_{multiTrait}^{p2}=max(H_{1}^{p2}, H_{2}^{p2}, \ldots, H_{n}^{p2}$)

…

$H_{multiTrait}^{pm}=max(H_{1}^{pm}, H_{2}^{pm}, \ldots, H_{n}^{pm})$,

$H_{multiTrait}^{residual}=max(H_{1}^{residual}, H_{2}^{residual}, \ldots, H_{n}^{residual})$,

where $max()$ means to return the vertex-wise maximum value among the n projected heritability maps. $H_{multiTrait}^{pi}$ represents the combined heritability maps of multiple AI-derived traits within $I_{i}$ component, and $H_{multiTrait}^{residual}$ represents the combined heritability maps of multiple AI-derived traits within the residual component.

6) Combine multiple independent components. Because independent components are statistically uncorrelated by construction, their contributions can be combined via an *L2*-norm aggregation.:

$$H_{multiTrait}^{multiIC}=\sqrt[2]{{H_{multiTrait}^{p1}}^{2}+{H_{multiTrait}^{p2}}^{2}+\ldots+{H_{multiTrait}^{pm}}^{2}{{+ H}_{multiTrait}^{residual}}^{2}}$$

Where $H_{multiTrait}^{multiIC}$ is the final heritability heatmap.

In step (5), we used the vertex-wise maximum across AI-derived traits when combining projected heritability maps within each independent component. This choice treats AI-derived traits as providing **separate but complementary discoveries**, each corresponding to an independently identified heritable facial trait, and as offering spatially localized evidence regarding where heritable variation manifests on the face. For example, if one AI-derived trait attributes a high heritable contribution to a specific facial region (e.g., nose) while another attributes heritability to a different region (e.g., chin), averaging would artificially dilute the localized signal in each region despite both being valid. The maximum operation therefore captures the “envelope” of heritable signal across AI-derived traits within a component.

In contrast, in Step (6), independent components represent **orthogonal latent axes of variation derived from the same set of AI-derived traits**, rather than separate discoveries. Assuming statistical independence across the components, their contributions are therefore combined using an *L2*-norm averaging to reflect the joint magnitude of effects across orthogonal dimensions.

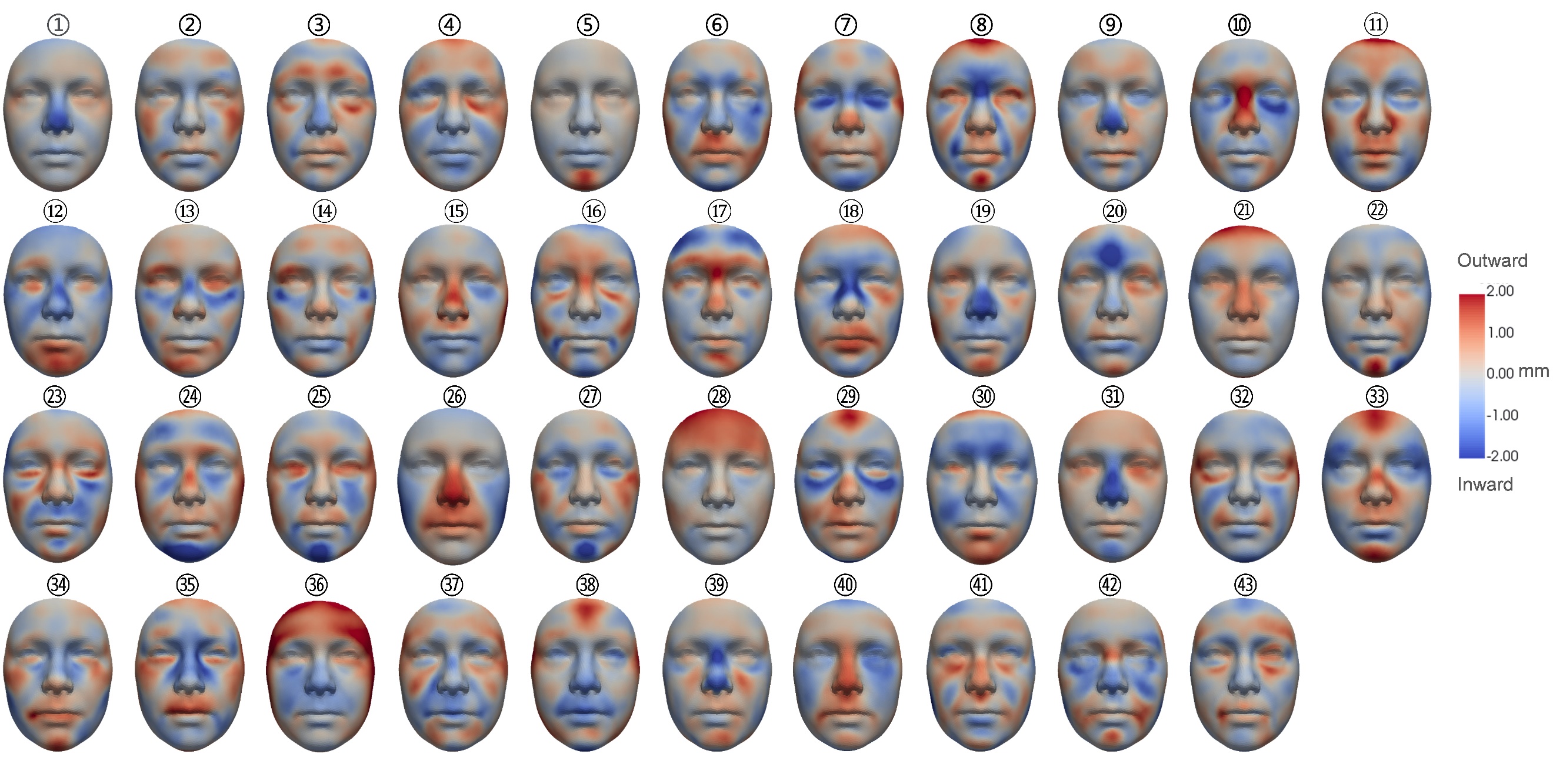

Figure S6: AI-based facial visualisation of the association effects of all 43 genetic loci with genome-wide significant associations in the discovery C-GWAS of both cohorts and all 200 traits combined.

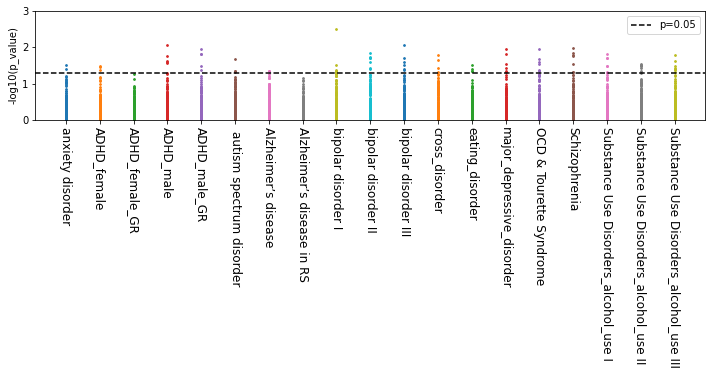

Figure S7: Genetic correlations between 200 AI-derived facial traits and diseases. For each column, one dot represents one trait.

Table S1: Genome-wide significant genetic loci from C-GWAS of both cohorts and all 200 AI-derived traits, with reported replication results from previously published independent European datasets (N = 8,246)^5^.

| index | Novel | SNPID | CHR | BP | Candidate genes | C-GWAS p-value | A1 | A2 | MAF | Replication p-value |
| --- | --- | --- | --- | --- | --- | --- | --- | --- | --- | --- |
| 1 | No | rs4648379 | 1 | 3261516 | *PRDM16* | 2.80E-08 | T | C | 0.2879 | 2.78e-19 |
| 2 | No | rs10923738 | 1 | 119594663 | *WARS2* | 1.86E-15 | A | G | 0.1627 | 1.42E-36 |
| 3 | Yes | rs12026445 | 1 | 161714087 | *FCRLA* | 3.11E-09 | G | A | 0.06937 | 2.95E-01 |
| 4 | Yes | rs9287125 | 1 | 192625360 | *RGS13* | 3.63E-08 | A | G | 0.06171 | 2.62E-01 |
| 5 | No | rs3122635 | 1 | 197324195 | *CRB1* | 3.44E-28 | T | G | 0.221 | 1.16E-68 |
| 6 | No | rs4832598 | 2 | 18702917 | *LOC105373454* | 1.98E-08 | A | G | 0.4728 | 5.77E-07 |
| 7 | Yes | rs6728062 | 2 | 37221985 | *HEATR5B* | 2.87E-08 | G | C | 0.1918 | 3.65E-01 |
| 8 | No | rs970797 | 2 | 177111819 | *MTX2/HOXD1/RPSAP25* | 3.15E-40 | T | G | 0.4485 | 1.13E-63 |
| 9 | No | rs260746 | 2 | 200114739 | NA | 3.17E-08 | C | A | 0.4826 | 3.24E-06 |
| 10 | No | rs752172 | 2 | 223092217 | *PAX3* | 6.18E-10 | A | G | 0.1401 | 2.98E-51 |
| 11 | No | rs76248430 | 3 | 25042638 | *RARB* | 3.78E-08 | A | C | 0.01222 | NA |
| 12 | Yes | rs115376369 | 4 | 55678447 | *LOC105377657* | 2.33E-08 | A | C | 0.04793 | 5.03E-01 |
| 13 | No | rs1021062 | 4 | 127969058 | NA | 4.06E-10 | T | C | 0.3356 | 1.53E-11 |
| 14 | No | rs35170094 | 4 | 128279531 | *LOC102724210* | 4.60E-08 | T | C | 0.3015 | 2.26E-05 |
| 15 | No | rs9995821 | 4 | 154828366 | *SFRP2/DCHS2/LOC101927947* | 1.21E-12 | C | T | 0.1969 | 9.30E-61 |
| 16 | No | rs1422446 | 5 | 128110085 | *LOC10537916*8 | 4.13E-10 | C | T | 0.3468 | 2.49E-12 |
| 17 | No | rs6555969 | 5 | 171128464 | *FGF18/SMIM23* | 3.10E-08 | T | C | 0.3212 | 2.33E-11 |
| 18 | No | rs9463083 | 6 | 45272062 | *SUPT3H* | 1.25E-11 | G | T | 0.4163 | 1.40E-24 |
| 19 | Yes | rs200459027 | 6 | 81725911 | NA | 4.37E-08 | G | GCTGGTA | 0.173 | 2.20E-01 |
| 20 | No | rs511490 | 6 | 133568151 | *EYA4* | 1.21E-15 | T | C | 0.2604 | 4.38E-38 |
| 21 | Yes | rs9496542 | 6 | 143569589 | *AIG1/LOC124901231* | 2.26E-08 | T | C | 0.3633 | 1.58E-03 |
| 22 | No | rs112159064 | 7 | 96155985 | *SEM1* | 3.16E-34 | G | GGGCTCT | 0.2922 | 3.30E-42 |
| 23 | No | rs142123729 | 7 | 101456740 | NA | 2.74E-08 | GT | G | 0.1249 | 5.01E-01 |
| 24 | No | rs12666947 | 7 | 120887544 | *CPED1/LOC124901735* | 1.38E-20 | T | C | 0.3443 | 1.39E-08 |
| 25 | No | rs34490323 | 9 | 16953117 | NA | 1.73E-10 | C | G | 0.421 | 1.31E-11 |
| 26 | Yes | rs201654567 | 10 | 62343275 | *ANK3/*  *LOC124902430* | 2.75E-08 | T | TG | 0.4962 | 1.08E-01 |
| 27 | No | rs242983 | 10 | 119276361 | *EMX2OS* | 1.51E-11 | A | G | 0.126 | 7.36E-18 |
| 28 | Yes | rs61877007 | 11 | 9996241 | *SBF2* | 1.08E-08 | C | T | 0.4034 | 8.41E-02 |
| 29 | No | rs6484256 | 11 | 10436441 | *CAND1.11* | 4.22E-08 | G | A | 0.203 | 5.86e-01 |
| 30 | Yes | rs113523945 | 11 | 75805724 | *UVRAG* | 1.27E-08 | G | A | 0.02015 | 3.69E-02 |
| 31 | No | rs35167392 | 12 | 28517664 | *CCDC91* | 9.02E-09 | T | TA | 0.4424 | 9.16E-08 |
| 32 | No | rs11175770 | 12 | 65851829 | *MSRB3* | 3.08E-12 | T | C | 0.4428 | 5.57E-16 |
| 33 | No | rs67713763 | 12 | 66377279 | NA | 8.50E-15 | A | T | 0.2102 | 2.87E-10 |
| 34 | No | rs6539808 | 12 | 84360625 | *LOC107984536* | 9.07E-09 | C | G | 0.389 | 2.59E-08 |
| 35 | No | rs11609649 | 12 | 85576492 | *LRRIQ1/ALX1* | 3.15E-13 | A | G | 0.172 | 5.35E-26 |
| 36 | Yes | rs2325624 | 13 | 74934563 | *LOC105370259* | 3.93E-08 | G | A | 0.07915 | 6.27E-01 |
| 37 | Yes | rs138028117 | 15 | 76883294 | *SCAPER* | 4.27E-08 | A | C | 0.0223 | 9.46E-01 |
| 38 | No | rs12940346 | 17 | 68075324 | *KCNJ16* | 5.23E-13 | C | T | 0.3522 | 1.70E-05 |
| 39 | No | rs11656479 | 17 | 69134717 | *CASC17* | 7.85E-17 | C | T | 0.3641 | 2.79E-38 |
| 40 | Yes | rs9949558 | 18 | 54914593 | *BOD1L2* | 2.16E-08 | G | A | 0.2669 | 8.94E-01 |
| 41 | No | rs6137476 | 20 | 21897656 | NA | 2.85E-08 | A | G | 0.4338 | 6.04E-16 |
| 42 | No | rs76855834 | 20 | 38001524 | NA | 3.11E-12 | T | C | 0.474 | 9.00E-20 |
| 43 | No | rs2834748 | 21 | 36453535 | NA | 2.75E-08 | G | A | 0.4748 | 1.16E-05 |

Table S2: Previously reported face-associated SNPs from 440 genetic loci.

| Gene | SNPID | CHR | BP | Loci number | PMID | Study | P |
| --- | --- | --- | --- | --- | --- | --- | --- |
| PRDM16 | rs76244841 | 1 | 2775953 | 1 | 33288918 | 2021 White et al. [5] | 1.48E-08 |
| PRDM16 | rs2377039 | 1 | 2803806 | 1 | 35393595 | 2022 Zhang et al. [82] | 2.52E-10 |
| PRDM16 | rs10797388 | 1 | 3252781 | 1 | 36539420 | 2022 Xiong et al. [6] | 9.38E-08 |
| PRDM16 | rs36192537 | 1 | 3253241 | 1 | 33547071 | 2021 Bonfante et al. [38] | 3.36E-11 |
| PRDM16 | rs7516137 | 1 | 3253839 | 1 | 35393595 | 2022 Zhang et al. [82] | 9.75E-29 |
| PRDM16 | rs1572037 | 1 | 3254369 | 1 | 33288918 | 2021 White et al. [5] | 2.64E-22 |
| PRDM16 | rs4648379 | 1 | 3261516 | 1 | 23028347 | 2012 Liu et al.  [83] | 1.13E-08 |
| PRDM16 | rs4648379 | 1 | 3261516 | 1 | 33983923 | 2021 Hoskens et al.  [37] | 8.52E-09 |
| CASZ1 | rs143353512 | 1 | 10733079 | 2 | 31763980 | 2019 Xiong et al.  [9] | 6.44E-09 |
| CASZ1 | rs143353512 | 1 | 10733079 | 2 | 36539420 | 2022 Xiong et al.  [6] | 2.69E-07 |
| CASZ1 | rs113365915 | 1 | 11002956 | 2 | 35393595 | 2022 Zhang et al.  [82] | 1.71E-10 |
| FBLIM1 | rs71631750 | 1 | 16083898 | 3 | 35393595 | 2022 Zhang et al.  [82] | 3.65E-08 |
| ARHGEF19 | rs200243292 | 1 | 16528473 | 3 | 31763980 | 2019 Xiong et al.  [9] | 1.46E-08 |
| PAX7 | rs4920524 | 1 | 18978372 | 4 | 33983923 | 2021 Hoskens et al.  [37] | 2.36E-08 |
| PAX7 | rs742071 | 1 | 18979874 | 4 | 33288918 | 2021 White et al.  [5] | 1.07E-15 |
| PAX7 | rs12410669 | 1 | 18990218 | 4 | 33547071 | 2021 Bonfante et al.  [38] | 3.58E-09 |
| PAX7 | rs111467622 | 1 | 18996519 | 4 | 35393595 | 2022 Zhang et al.  [82] | 1.53E-25 |
| CAPZB | rs4912082 | 1 | 19716796 | 5 | 33288918 | 2021 White et al.  [5] | 3.22E-08 |
| CAPZB | rs4912094 | 1 | 19762466 | 5 | 35393595 | 2022 Zhang et al.  [82] | 1.26E-09 |
| EIF4G3 | rs11462489 | 1 | 21310047 | 6 | 36539420 | 2022 Xiong et al.  [6] | 2.39E-06 |
| CDC42 | rs55703910 | 1 | 22785206 | 7 | 33983923 | 2021 Hoskens et al.  [37] | 3.80E-08 |
| EPHB2 | rs75125839 | 1 | 23309666 | 8 | 33983923 | 2021 Hoskens et al.  [37] | 3.19E-08 |
| HNRNPR | rs139879053 | 1 | 23639799 | 8 | 27560698 | 2016 Cole et al.  [81] | 2.73E-08 |
| MATN1 | rs16834081 | 1 | 31292641 | 9 | 33288918 | 2021 White et al.  [5] | 2.35E-09 |
| ZSCAN20 | rs16835519 | 1 | 33966753 | 10 | 33983923 | 2021 Hoskens et al.  [37] | 8.59E-09 |
| - | esv2656640 | 1 | 38602524 | 11 | 33983923 | 2021 Hoskens et al.  [37] | 4.48E-08 |
| MACF1 | rs1021829 | 1 | 39591141 | 12 | 33983923 | 2021 Hoskens et al.  [37] | 3.92E-08 |
| GLIS1 | rs72906949 | 1 | 53970693 | 13 | 35393595 | 2022 Zhang et al.  [82] | 4.55E-09 |
| GLIS1 | rs3887651 | 1 | 53999509 | 13 | 35393595 | 2022 Zhang et al.  [82] | 1.32E-18 |
| GLIS1 | rs12089978 | 1 | 54045180 | 13 | 33983923 | 2021 Hoskens et al.  [37] | 3.34E-08 |
| GLIS1 | rs2948053 | 1 | 54062273 | 13 | 35393595 | 2022 Zhang et al.  [82] | 3.04E-11 |
| GLIS1 | rs702489 | 1 | 54197688 | 13 | 27182965 | 2016 Pickrell et al.  [84] | 2.50E-09 |
| … | … | … | … | … | … | … | … |
| SPANXN1 | rs112437700 | 23 | 144270385 | 440 | 33983923 | 2021 Hoskens et al.  [37] | 4.60E-08 |

CHR: Chromosome

BP: Basepairposition

PMID: PubMedIdentifier.

Previously reported face-associated SNPs were selected from 20 studies, as shown in columns: PMID and study.

The reported p-value of SNPs and their candidate genes are indicated in columns: P and Gene.

We clumped SNPs into 440 loci separated by at least 500kb, shown in column: Loci number.

The full table, which includes 941 rows, can be found online: https://docs.google.com/spreadsheets/d/16lf8JYe6IyhNMxddmRPRwMnKqBFBXgsE

Table S3: Heritability of the 200 AI-derived traits estimated via LDSC-based heritability analysis on the single-trait GWAS summary after meta-analysis of the two cohorts.

| **AI-derived trait** | **h2** | **FDR-corrected p value** |
| --- | --- | --- |
| **158** | 0.3448 | 4.21e-06 |
| **83** | 0.2974 | 4.25e-04 |
| **23** | 0.2795 | 5.95e-04 |
| **79** | 0.2729 | 5.95e-04 |
| **145** | 0.2721 | 5.95e-04 |
| **21** | 0.2629 | 9.63e-04 |
| **192** | 0.2627 | 5.95e-04 |
| **150** | 0.2590 | 5.95e-04 |
| **172** | 0.2529 | 2.63e-03 |
| **…** | … | … |
| **135** | 0.126 | 4.81e-02 |

In total, 90 traits were found FDR-significant in the heritability analysis.

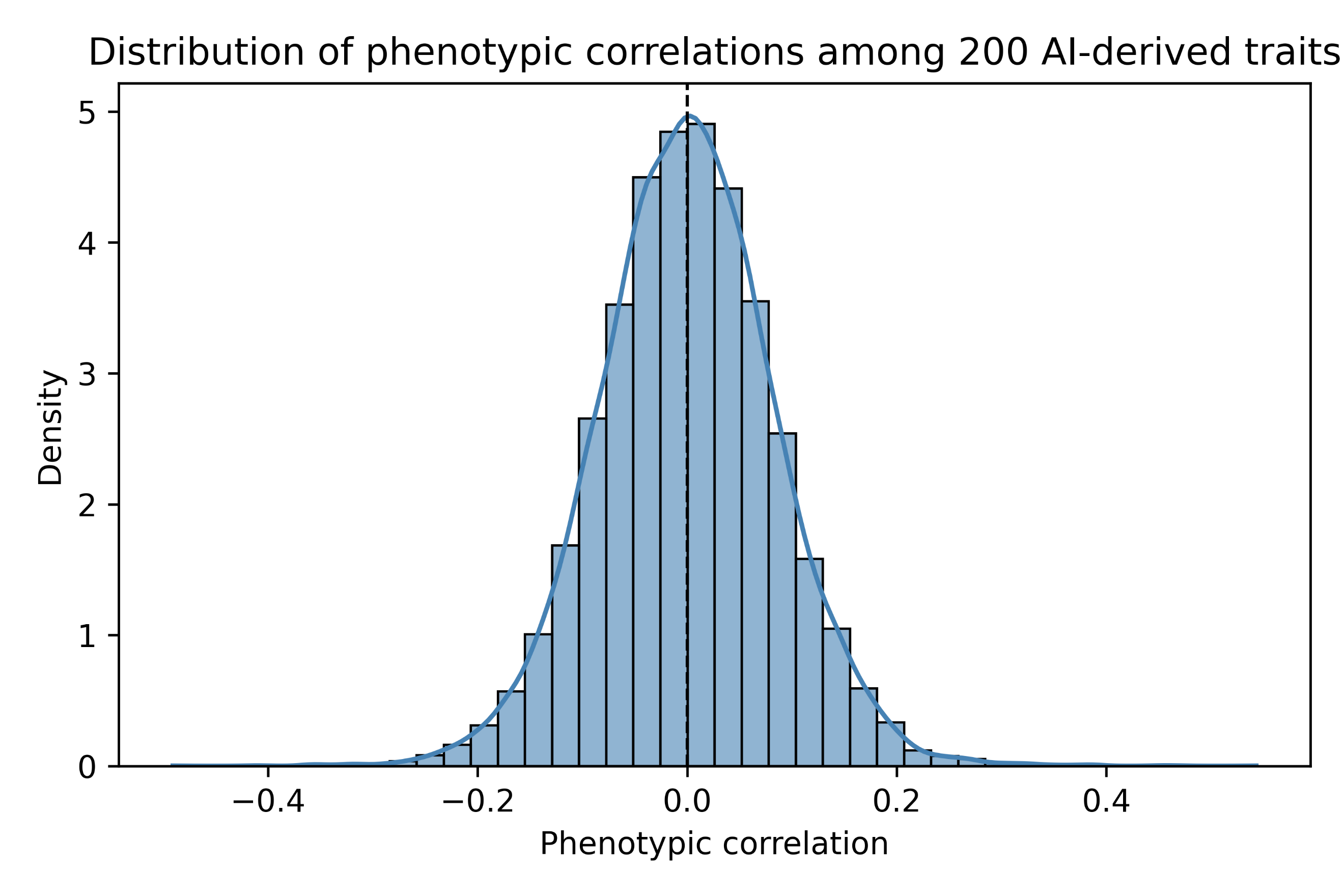

Figure S8: Histogram distribution of phenotypic correlations among 200 AI-derived traits.

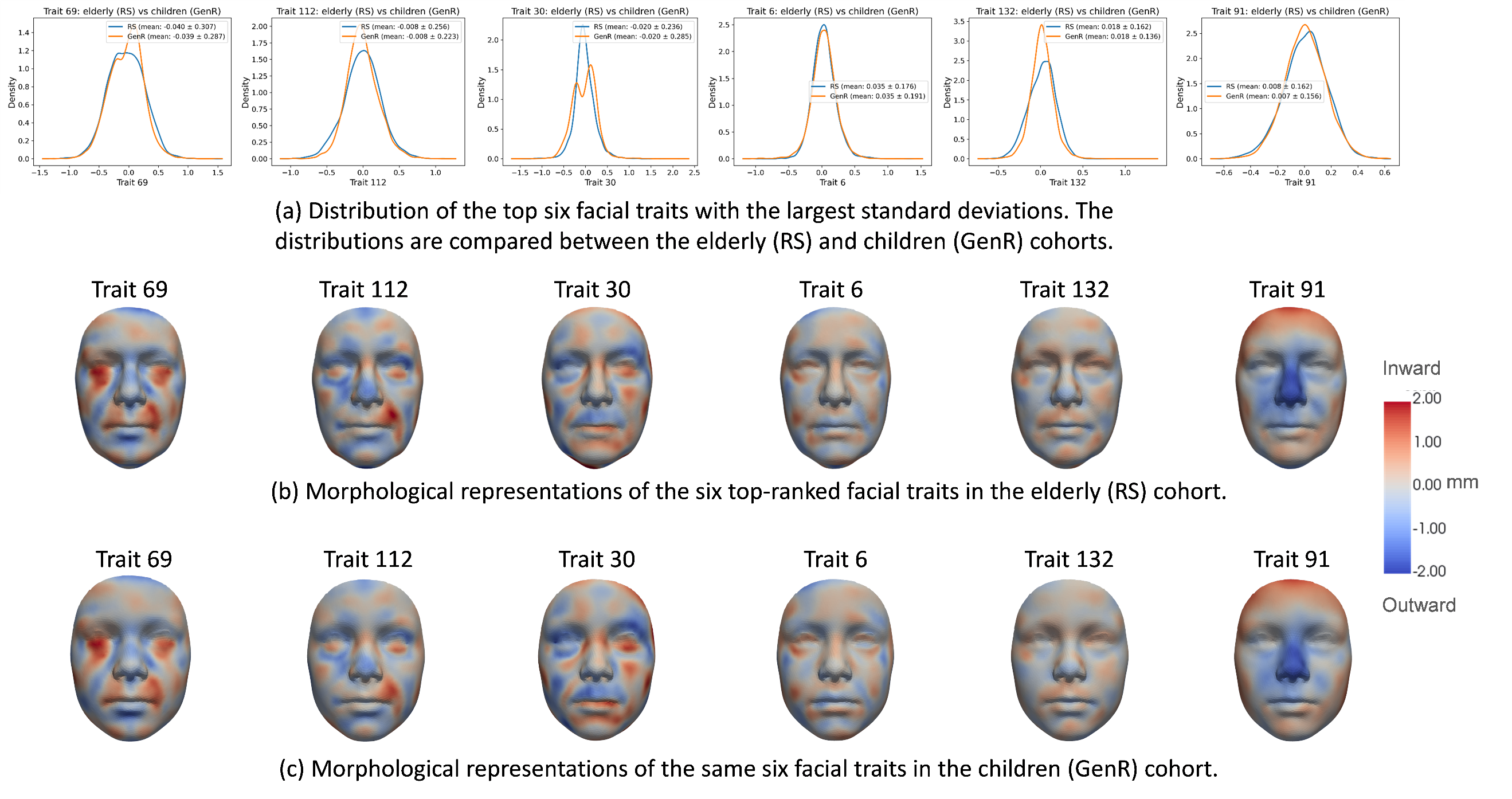

Figure S9: Alignment of AI-derived facial traits across cohorts based on the federated learning (FL) model. Representative facial traits derived from the FL-trained model are visualized for both the Rotterdam Study (RS) and Generation R (GenR) cohorts. The corresponding traits exhibit consistent facial morphological patterns across cohorts, indicating that the FL framework aligns the learned latent representations and enables comparable phenotyping across datasets without sharing individual-level image data.

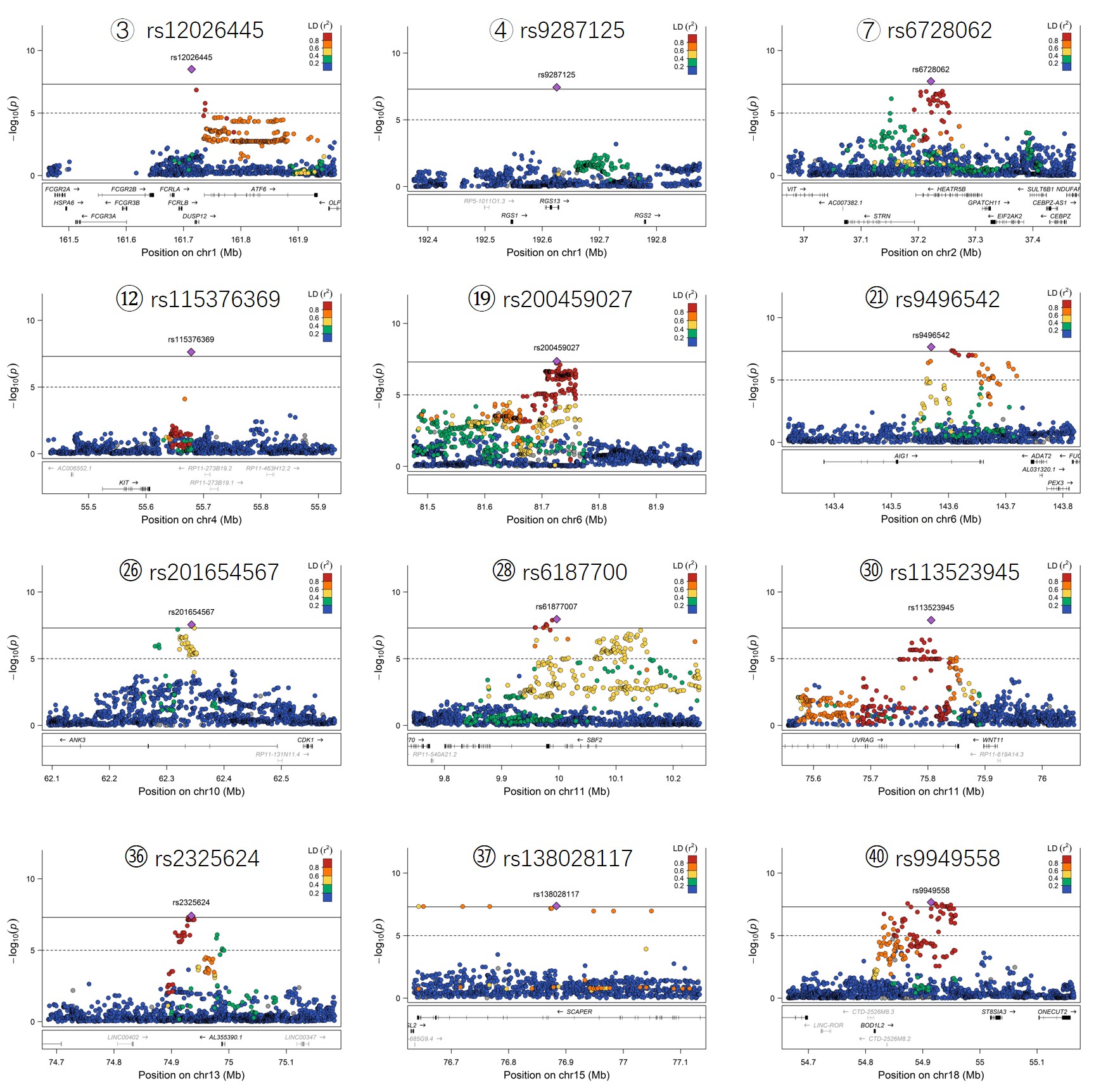

Figure S10: Regional association plots of the 12 novel loci identified in this study.
